# Membrane constriction and thinning by sequential ESCRT-III polymerization

**DOI:** 10.1101/798181

**Authors:** Henry C. Nguyen, Nathaniel Talledge, John McCullough, Abhimanyu Sharma, Frank R. Moss, Janet H. Iwasa, Michael D. Vershinin, Wesley I. Sundquist, Adam Frost

## Abstract

The Endosomal Sorting Complexes Required for Transport (ESCRTs) mediate diverse membrane remodeling events. These activities typically require ESCRT-III proteins to stabilize negatively-curved membranes, although recent work has indicated that certain ESCRT-IIIs also participate in positive-curvature membrane shaping reactions. ESCRT-IIIs polymerize into membrane-binding filaments, but the structural basis for negative versus positive membrane curvature shaping by these proteins remains poorly understood. To learn how ESCRT-IIIs shape membranes, we determined structures of human membrane-bound CHMP1B-only, membrane-bound CHMP1B+IST1, and IST1-only filaments by electron cryomicroscopy. Our structures show how CHMP1B first polymerizes into a single-stranded helical filament, shaping membranes into moderate-curvature tubules. Subsequently, IST1 assembles a second strand upon the CHMP1B filament, further constricting the membrane tube and reducing its diameter nearly to the fission point. Each step of constriction, moreover, thins the underlying bilayer and lowers the barrier to membrane fission. Together, our structures reveal how a two-component, sequential polymerization mechanism drives membrane tubulation, tube constriction, and bilayer thinning.

## Introduction

The Endosomal Sorting Complexes Required for Transport (ESCRT) belong to an evolutionarily conserved pathway that mediates membrane remodeling and fission events throughout the cell. The ESCRT machinery comprises staged complexes, including the early-acting ALIX, ESCRT-I, and -II factors and the late-acting ESCRT-III factors and VPS4 family of AAA+ ATPases. Early-acting factors bind to site-specific adaptors and then recruit the late-acting factors that constrict and sever the target membrane. First discovered for their role in the formation of multivesicular bodies (MVBs), ESCRT proteins serve essential functions in an expanding range of cellular processes. Beyond MVBs, these processes include: cytokinetic abscission; egress of enveloped viruses; sealing holes in nuclear, endosomal, and plasma membranes (*1–9*); and in peroxisome biogenesis and function (*10, 11*). ESCRT-III proteins primarily shape negatively-curved membranes, such as the necks of budding viruses or intralumenal vesicles, but we and others have shown that some ESCRT-III proteins can also stabilize positively-curved membranes (*12–15*). Despite their importance to the cell, the mechanisms that govern how ESCRT-III proteins assemble and catalyze membrane remodeling reactions—of either positive or negative membrane curvature—remain unclear.

Humans have 12 different ESCRT-III proteins that share a conserved secondary structure core, including helices α1-α5. X-ray crystal structures of IST1 and CHMP3 revealed how these helices fold into a compact conformation referred to as a “closed” state (*16–18*). Other ESCRT-III proteins can adopt more elongated “open” states that can also polymerize (*16, 19–21*). Structures of such open, elongated and assembled states are available for human CHMP1B (*14*), *S. cerevisiae* Snf7 (*22*), and *D. melanogaster* Shrub (*23*). ESCRT-III polymerization may be regulated by reversible switching between closed and open conformations. Such conformational transitions could be regulated by protein-protein interactions with nucleating factors like the early-acting ALIX or ESCRT-II factors, by membrane curvature (*24*), or by post-translation modifications, such as ubiquitination, which can sterically hinder membrane-binding or polymerization (*25*).

We previously reported a ∼4 Å resolution electron cryomicroscopy (cryoEM) structure of a helical filament containing CHMP1B and IST1 (*14*). This copolymer structure was surprising as it consisted of two distinct strands: an inner strand of CHMP1B in an open conformation and an outer strand of IST1 in a closed conformation. The IST1 strand was tightly associated with the CHMP1B strand with 1:1 stoichiometry. The lumenal cavity of the copolymer was strongly positively charged and capable of shaping negatively charged membranes into positive-curvature membrane tubes in vitro and in vivo. Consistently, studies in living cells have shown that CHMP1B and IST1 co-localize with the VPS4 family member SPASTIN along the positive curvature surfaces of endosomal tubules and contact sites between lipid droplets and peroxisomes (*10, 14, 15, 26*). To better understand these new properties and roles for CHMP1B and IST1, and the still unknown structural mechanism by which any ESCRT-III protein interacts with lipid bilayers, we sought to understand how CHMP1B and IST1 work together to bind and constrict membranes.

## Results

### Structure of the membrane-bound CHMP1B filament

To learn how CHMP1B and IST1 remodel positively curved membranes, we sought to capture stable membrane-bound polymers composed of these proteins. We previously showed that incubating liposomes with CHMP1B led to formation of membrane tubules coated with protein filaments (*14*). To increase the yield and stability of these membrane tubules, we optimized the lipid composition and found that CHMP1B could remodel a variety of different liposome compositions into tubules. Two factors, in particular, enhanced the prevalence and stability of membrane-bound filaments for cryoEM analysis (Figure 1A and 1D): 1) incorporation of polyunsaturated lipids (16:0-22:6 phosphocholine, SDPC) to increase membrane malleability (*27, 28*); and 2) increasing the concentration of negatively charged phospholipids such as PI(3)P (phosphatidylinositol 3-phosphate), PI(3, 5)P_2_, or PI(4, 5)P_2_ to complement the highly basic charge of the CHMP1B lumen (*14*). For our in-depth studies, we settled on liposomes containing 58 mol% SDPC / 18 mol% POPS / 18 mol% cholesterol / 6 mol% PI(3, 5)P_2_.

**Figure 1.**
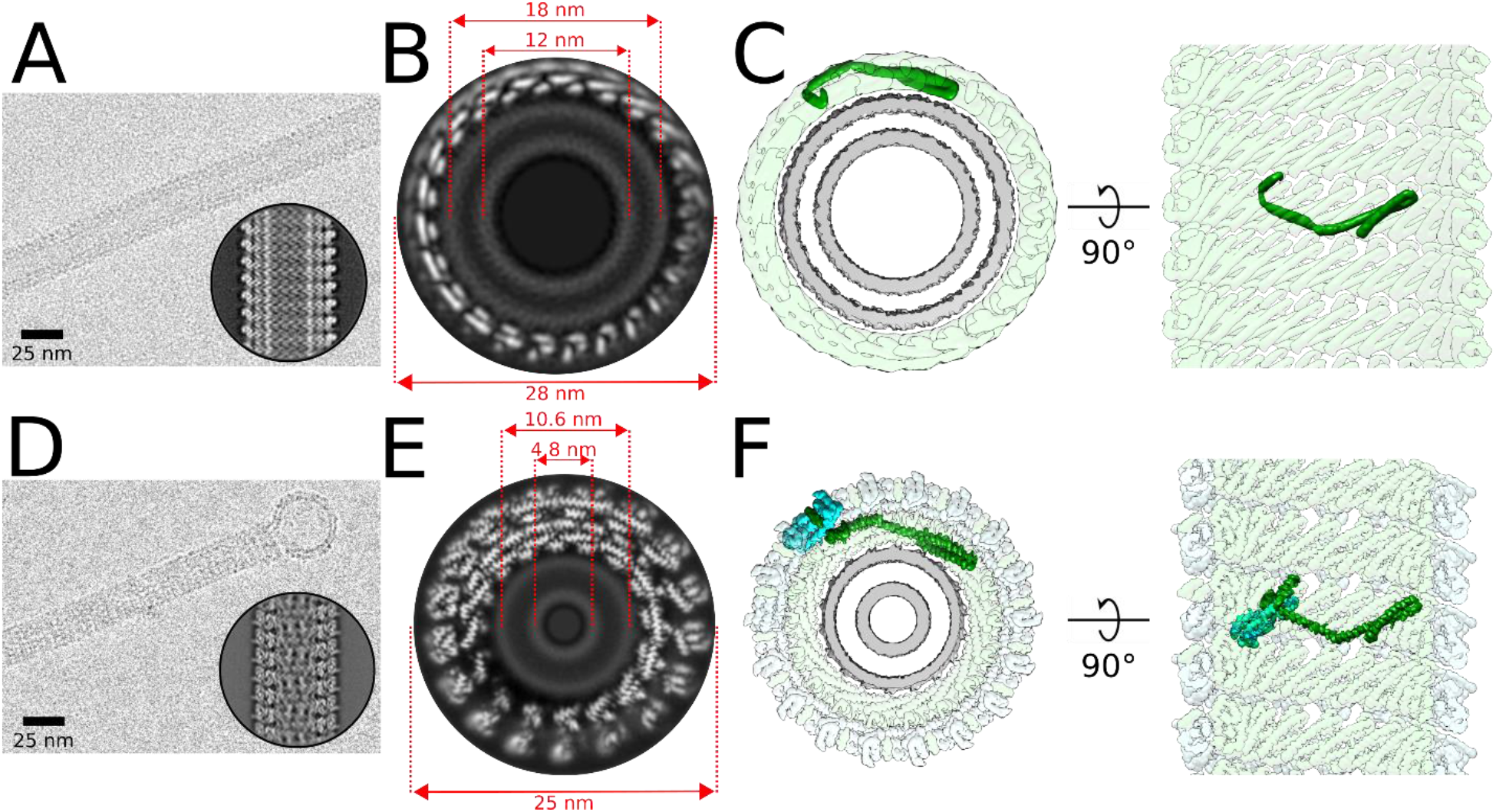
CHMP1B and IST1 sequentially constrict membrane tubes. (**A**) CryoEM micrograph of a membrane-bound CHMP1B tubule. Scale bar: 25 nm. Inset, representative 2D class average. (**B**) A grey-scale slice looking down the helical axis of the 3D cryoEM reconstruction of the membrane-bound CHMP1B filament. Diameters of the entire tube and the membrane leaflet peak-to-peak distances are annotated. (**C**) *Left*, surface representation of the same end-on view as in B down the helical axis. CHMP1B (green) coats the exterior of the membrane bilayer (grey). A CHMP1B protomer is highlighted in dark green. *Right*, internal view looking outward from the surface of the membrane. (**D**) IST1 further constricts the CHMP1B-membrane filament nearly to the hemifission point. CryoEM micrograph of a membrane-bound CHMP1B+IST1 filament with a vesicle protruding from the end. Scale bar: 25 nm. Inset, representative 2D average. (**E**) A grey-scale slice looking down the helical axis of the 3D cryoEM reconstruction of the membrane-bound, right-handed CHMP1B+IST1 filament. Diameters of the entire tube and membrane leaflet peak-to-peak distances are annotated. (**F**) *Left*, surface representation of the same end-on view as in E down the helical axis. (*Right*), internal view looking outward from the surface of the membrane. IST1 protomers (cyan) binds to the exterior of CHMP1B (green), leading to constriction of the membrane (grey). IST1 and CHMP1B promoters are highlighted in dark cyan and dark green, respectively.

**Figure 1—figure supplement 1.**
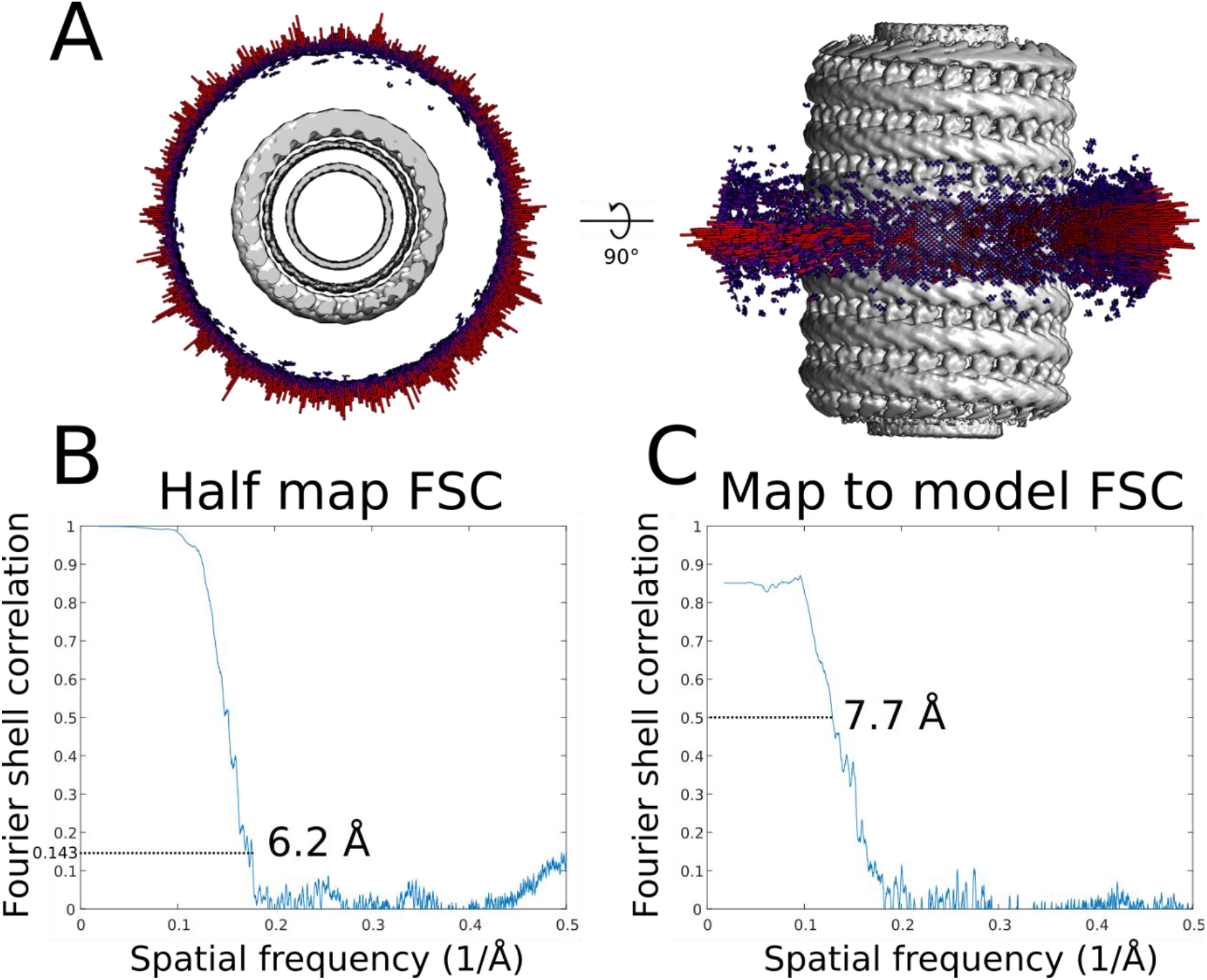
CryoEM validation of membrane-bound CHMP1B-only filament. (**A**) Angular distribution of the membrane-bound CHMP1B filament. (**B**) Half map Fourier shell correlation (FSC) of the membrane-bound CHMP1B filament. (**C**) Map to model FSC of the membrane-bound CHMP1B filament.

**Figure 1—figure supplement 2.**
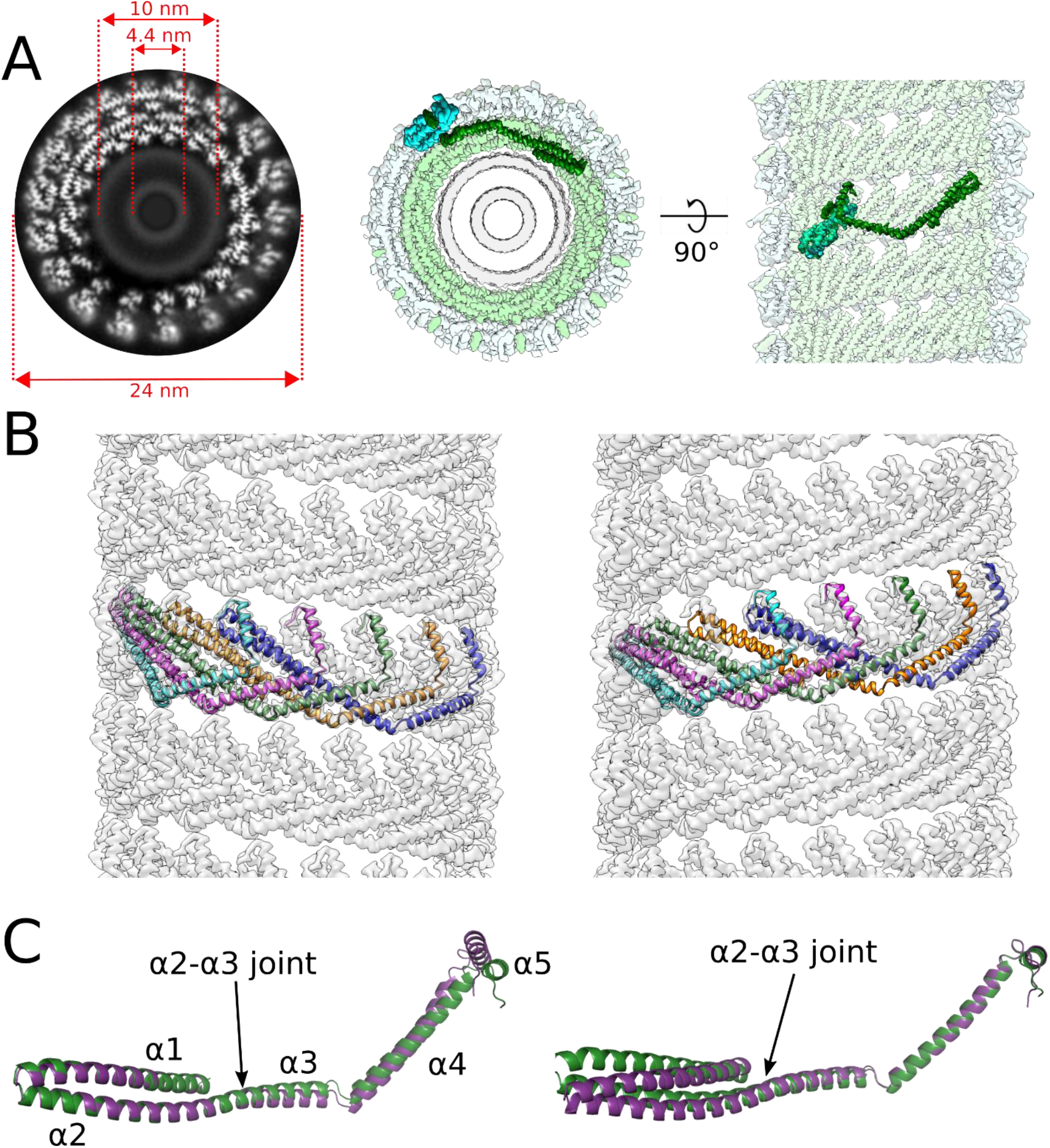
CryoEM reconstruction of the membrane-bound CHMP1B+IST1 filament at higher curvature and comparison of left- and right-handed CHMP1B+IST1 filaments. (**A**) CryoEM 3D reconstruction of the membrane-bound left-handed CHMP1B+IST1 filament. End-on view down the helical axis in grey-scale (*left*) or colored (*middle*). *Right*, internal view looking outward from the membrane surface along the helical axis. IST1 protomers (cyan) bind to the exterior of CHMP1B (green), leading to constriction of the membrane (grey). IST1 and CHMP1B promoters are highlighted in dark cyan and green, respectively. Diameters of the entire tube and membrane leaflet peak-to-peak distances are annotated. (**B**) Electron density maps of CHMP1B from the left-handed (*left*) or right-handed (*right*) membrane-bound CHMP1B+IST1 filaments. Five copies of CHMP1B are shown as ribbons. (**C**) Superposition of a CHMP1B protomer from the left-handed (purple) and right-handed (green) CHMP1B+IST1 filaments aligned to the CHMP1B N-terminal α1-α2 helices (*left*) or C-terminal α4-α5 helices (*right*).

**Figure 1—figure supplement 3.**
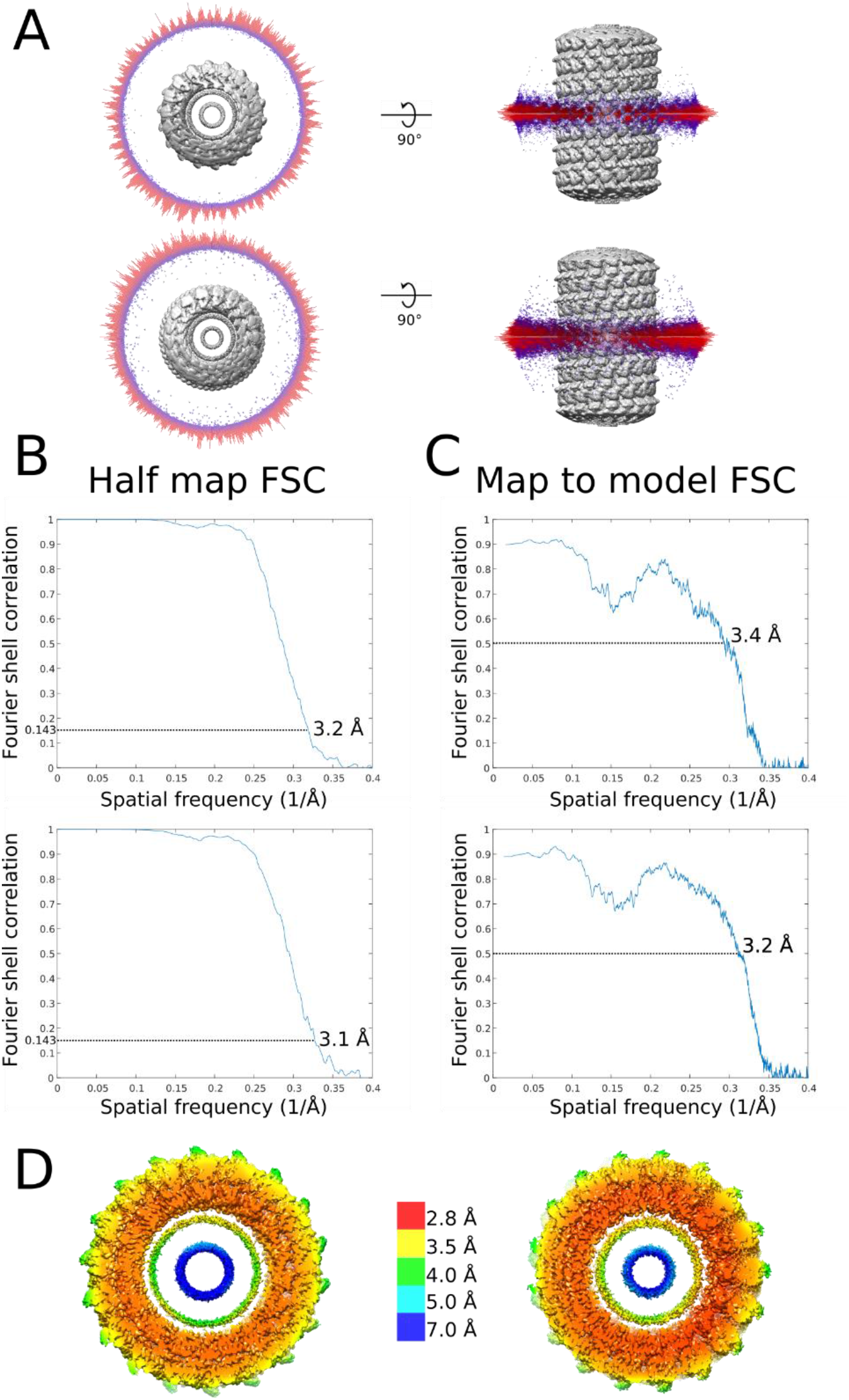
Local resolution estimates and cryoEM validation of membrane-bound CHMP1B+IST1 filaments. (**A**) Angular distribution of right-handed (*top*) and left-handed (*bottom*) membrane-bound CHMP1B+IST1 filaments. (**B**) Half map FSCs of right-handed (*top*) and left-handed (*bottom*) CHMP1B+IST1 filaments. (**C**) Map to model FSCs right-handed (*top*) and left-handed (*bottom*) CHMP1B+IST1 filaments. (**D**) Local resolution estimates of right-handed (*left*) and left-handed (*right*) CHMP1B+IST1 filaments.

From this optimized lipid mixture, we first analyzed CHMP1B-only membrane tubes and found that they exhibited a range of diameters (∼26-30 nm). This heterogeneity precluded high-resolution studies, but by sorting the tubes based on diameter, we were able to reconstruct a 28 nm diameter tube to ∼6 Å resolution. This reconstruction unambiguously showed the open-state conformation of CHMP1B and the interconnected network of protomers within the single-stranded, right-handed filament (Figure 1B–1C, Figure 1—figure supplement 1, Tables 1-3). Upon membrane tubulation, the membrane tube diameter, the distance between outer leaflet phosphate headgroups, narrows from >50 nm in the starting, spherical liposomes down to ∼18 nm in the CHMP1B-constricted cylindrical state. The inner leaflet headgroups are separated by ∼12 nm. Thus, the energy of CHMP1B self-assembly upon the membrane is sufficient to remodel low-curvature membrane spheres into moderate-curvature membrane cylinders.

**Table 1.** CryoEM data collection parameters.

| Dataset | CHMP1B-only | CHMP1B+IST1 | IST1<sub>NTD</sub><sup>R16E/K27E</sup> |
| --- | --- | --- | --- |
| Microscope | Titan Krios | Polara | TF20 |
| Energy filter (slit width, eV) | 20 | N/A | N/A |
| Voltage (kV) | 300 | 300 | 200 |
| C2 aperture (μm) | 70 | 30 | 70 |
| Objective aperture (μM) | 100 | 100 | 100 |
| Camera | K2 Summit | K2 Summit | K2 Summit |
| Detection mode | Super resolution | Super resolution | Super resolution |
| Pixel size (Å/Pixel) | 1.345 | 1.2156 | 1.234 |
| Total exposure (sec) | 10 | 8.0 | 8.0 |
| Frame rate (Frames/sec) | 0.2 | 0.2 | 0.2 |
| Dose rate (e<sup>-</sup>/Å<sup>2</sup>/frame) | 0.90 | 1.1 | 1.325 |
| Total dose (e<sup>-</sup>/Å<sup>2</sup>) | 45 | 44 | 53 |
| Defocus range (μm) | 0.23-1.91 | 0.46-2.97 | 1.04-3.22 |
| Mean defocus (μm) | 1.14 | 1.54 | 2.28 |

**Table 2.** CryoEM refinement parameters.

| Dataset | CHMP1B-only | CHMP1B+IST1<br>(right-handed) | CHMP1B+IST1<br>(left-handed) | IST1<sub>NTD</sub><sup>R16E/K27E</sup> |
| --- | --- | --- | --- | --- |
| Micrographs | 3993 | 9305 | 9305 | 279 |
| Segments in final map | 9,661 | 66,149 | 57,915 | 4,556 |
| Box size (Å<sup>3</sup>) | 324 | 352 | 352 | 320 |
| Resolution (Å) | 6.2 | 3.2 | 3.1 | 7.2 |
| Helical twist (°)* | +13.86 | +20.02 | -20.77 | +26.85 |
| Helical rise (Å) | 1.86 | 2.96 | 3.06 | 3.00 |
| Map sharpening B-factor (Å<sup>2</sup>) | -50 | -25 | -25 | -50 |
| Resolution (Å) | 6.2 | 3.2 | 3.1 | 7.2 |

\*denotes + for right-handed and - for left-handed twists

**Table 3.** Model validation statistics.

| Dataset | CHMP1B-only | CHMP1B+IST1<br>(right-handed) | CHMP1B+IST1<br>(left-handed) | IST1<sub>NTD</sub><sup>R16E/K27E</sup> |
| --- | --- | --- | --- | --- |
| Chains | 26 | 72 | 68 | 14 |
| Atoms | 32,890 | 99,216 | 93,500 | 19,432 |
| Protein residues | 4,238 | 12,852 | 12,104 | 2,534 |
| RMS bonds (Å) | 0.006 | 0.011 | 0.005 | 0.007 |
| RMS angles (°) | 0.816 | 0.828 | 0.863 | 1.15 |
| MolProbity score | 1.55 | 1.17 | 1.14 | 1.61 |
| Clashscore<br>(percentile relative to<br>PDB) | 10.73 (68<sup>th</sup>) | 3.82 (96<sup>th</sup>) | 7.29 (85<sup>th</sup>) | 5.48 (93<sup>rd</sup>) |
| Ramachandran<br>favored (%) | 98.76 | 98.58 | 98.86 | 95.53 |
| Ramachandran<br>allowed (%) | 1.24 | 1.42 | 1.14 | 4.47 |
| Ramachandran<br>outliers (%) | 0 | 0 | 0 | 0 |
| Rotamer outliers (%) | 0 | 0 | 0.34 | 0.74 |
| C-beta outliers | 0 | 0 | 0 | 0 |
| Mean B-factor | N/A* | 68 | 60 | N/A* |
| Minimum/maximum<br>B-factor | N/A* | 51/112 | 45/92 | N/A* |
| EMRinger score | N/A* | 1.05 | 1.41 | N/A* |

\*The resolutions of the maps are insufficient to determine these values

### Structures of left- and right-handed membrane-bound CHMP1B+IST1 filaments reveal highly constricted bilayers and filament flexibility

We next investigated the effect of sequentially adding IST1 to these filaments by cryoEM. Interestingly, from the same sample we were able to determine two 3D reconstructions of membrane-bound CHMP1B+IST1 filaments, corresponding to approximately equal populations of right- and left-handed helical filaments, to 3.2 Å and to 3.1 Å resolution, respectively (Figure 1E–1F, Figure 1—figure supplements 2–3, Tables 1–3). The right-handed CHMP1B+IST1 copolymer is a, one-start, double-stranded filament. The outer strand comprises IST1 in the closed conformation, and the inner strand comprises CHMP1B again in the open conformation (*14*). The reconstruction also reveals a continuous and highly constricted bilayer within the lumen. The overall outer diameter of the double-stranded filament is 25 nm, slightly narrower than the membrane-bound CHMP1B-only filament. However, due to the presence of two protein strands, the distance between outer leaflet phosphate headgroups is reduced to 10.6 nm, and the distance between inner leaflet headgroups is just 4.8 nm (Figure 1E). Thus, the sequential addition of IST1 was sufficient to drive constriction of the CHMP1B strand and the internal membrane, narrowing the lumenal inner leaflet diameter from 12 nm down to 4.8 nm.

**Figure 2.**
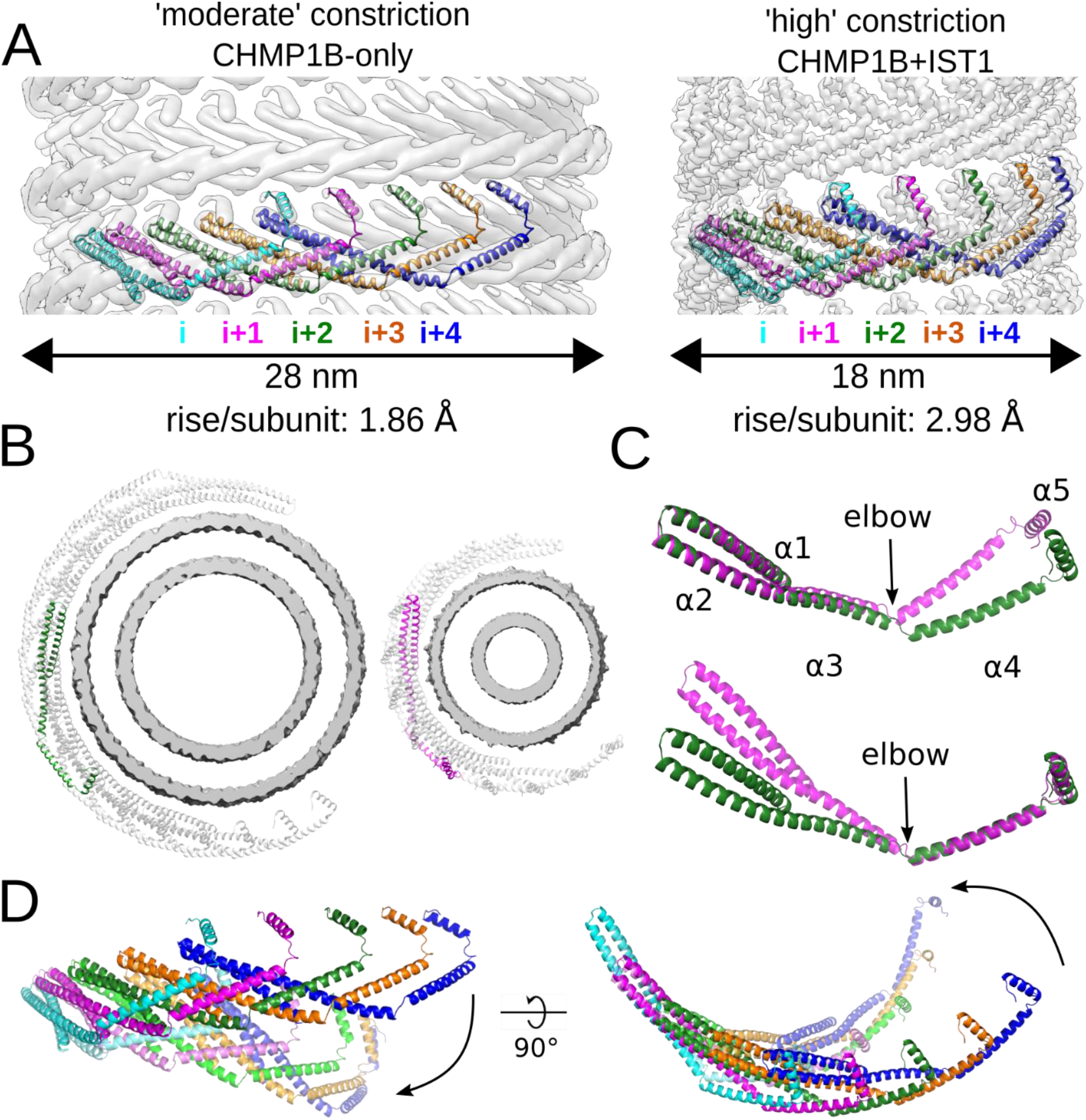
CHMP1B interlocks in the same arrangement in all structures and flexes at the α3-α4 elbow to accommodate different curvatures. (**A**) CryoEM density maps of CHMP1B from the membrane-bound CHMP1B (*left*) or right-handed CHMP1B+IST1 (*right*) filaments. Five interlocked copies of CHMP1B are shown as ribbons. The C-terminal helix α5 of the i protomer always engages helices α1-α2 of the i+4 protomer. The rise per subunit for each helical filament is denoted. (**B**) Comparison of arc curvatures of CHMP1B across the two filaments. Top-down views of half a turn of CHMP1B subunits are shown for either the CHMP1B (*left*) or CHMP1B+IST1 (*right*) membrane filaments. The membrane bilayers are shown in grey and the central promoters are shown in green and magenta for the respective filaments. (**C**) Superposition of a CHMP1B protomer from the CHMP1B (green) and CHMP1B+IST1 (magenta) filaments aligned to the CHMP1B N-terminal helices α1-α2 (top) or C-terminal helices α4-α5 (bottom). The biggest conformational change occurs at the elbow joint. (**D**) Superposition of 5 consecutive subunits (colored from left to right in cyan, magenta, green, orange, and blue) of CHMP1B from the CHMP1B (opaque) and the CHMP1B+IST1 (semi-transparent) filament. The respective first protomers from each are aligned as in (C).

**Figure 2—figure supplement 1.**
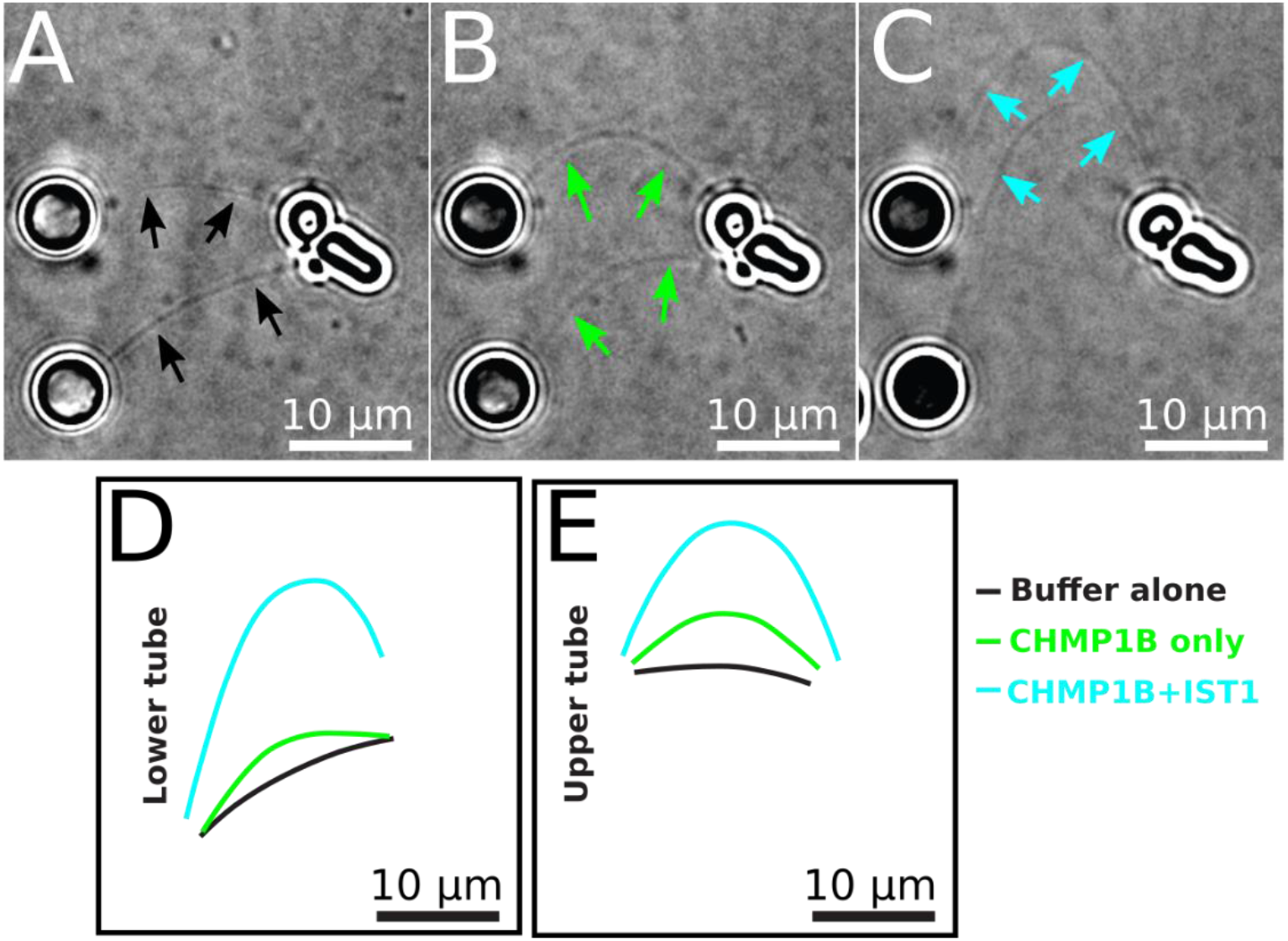
Real-time monitoring of CHMP1B and IST1 membrane constriction and elongation. (**A-C**) Still images representing deformation of two membrane tubes due to transverse flow of (A) buffer alone, (B) then 0.5 μM CHMP1B, (C) and a final addition of 0.5 μM IST1 is shown. Solid arrows in (A)-(C) highlight tubule locations. (**D-E**) Contours of lower (D) and upper (E) membrane tubes extracted from panels (A)-(C) showing the extension of the tubes upon addition of CHMP1B and IST1.

**Figure 2—movie supplement 1.**
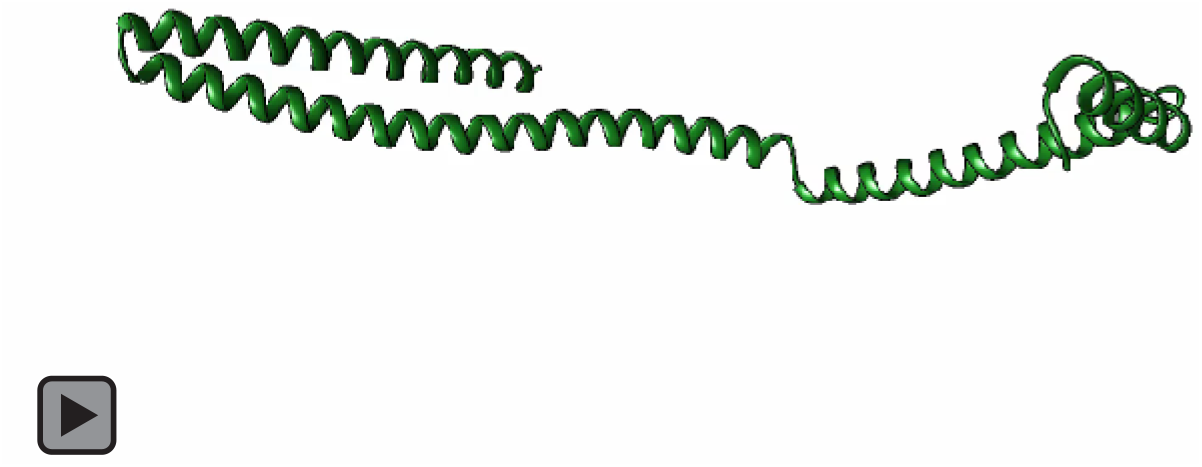
Elbow flexing of one CHMP1B subunit.

**Figure 2—movie supplement 2.**
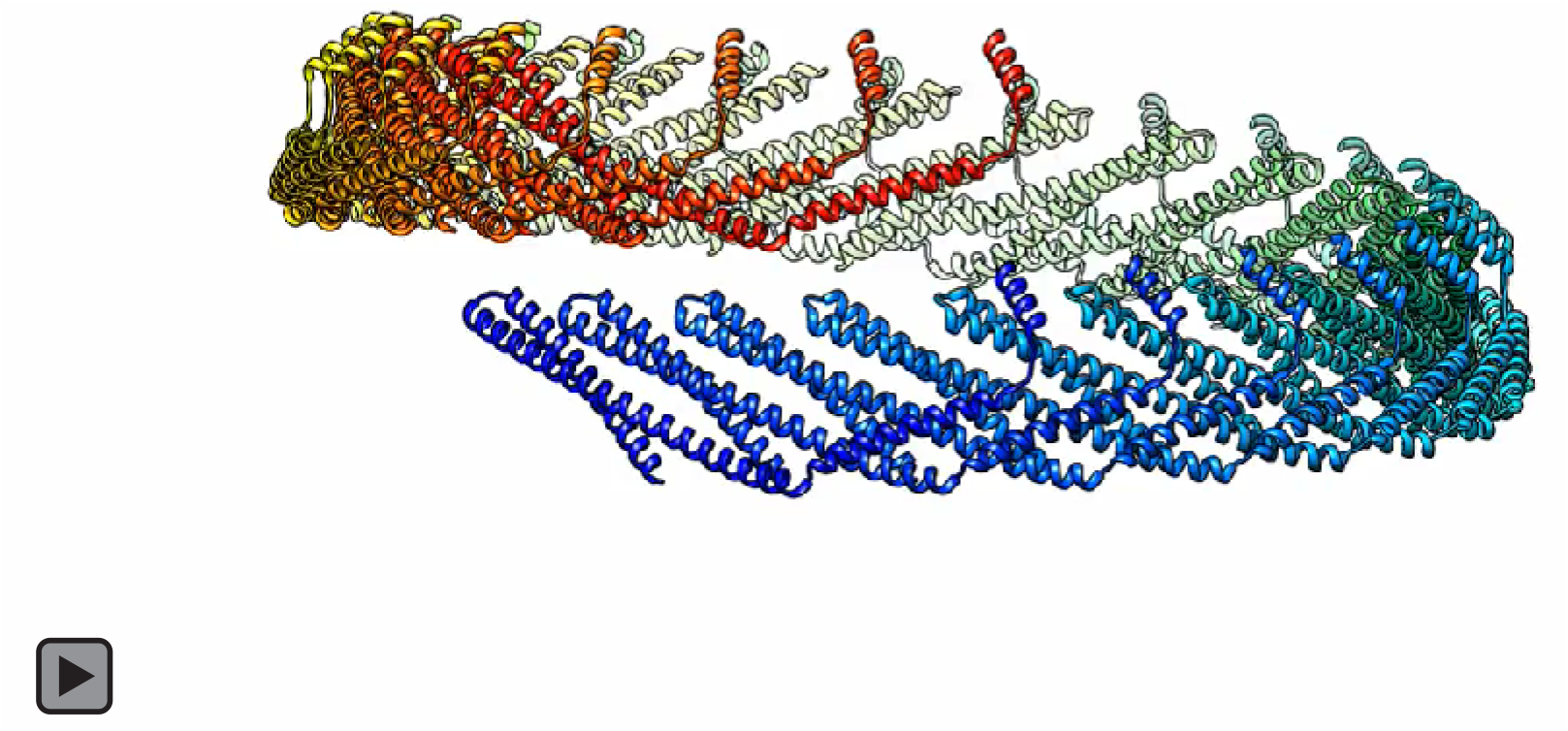
Flexing of a full turn of CHMP1B subunits from low to high constriction.

**Figure 2—movie supplement 3.**
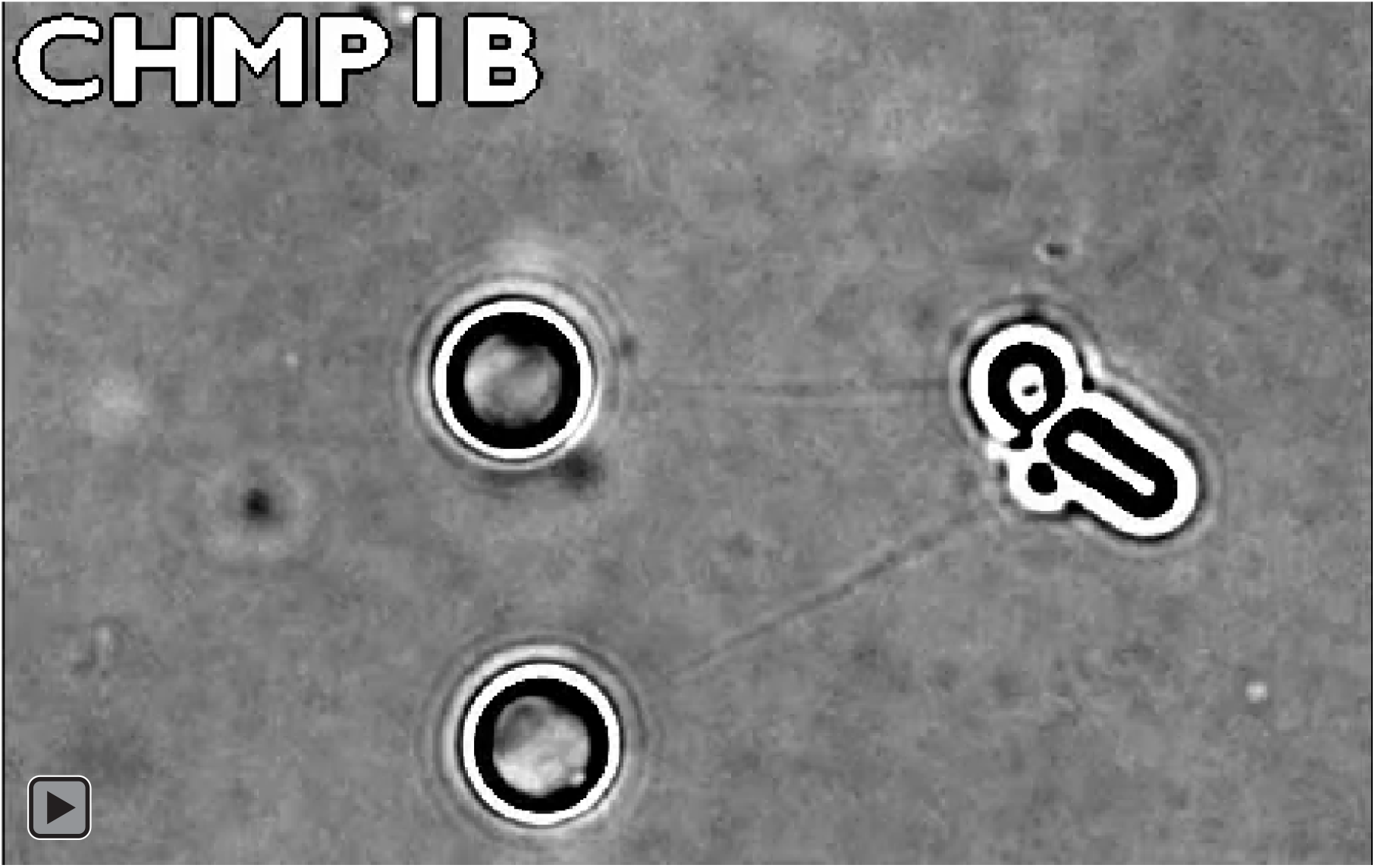
Real time recording of membrane tube elongation by CHMP1B and IST1.

**Figure 2—movie supplement 4.**
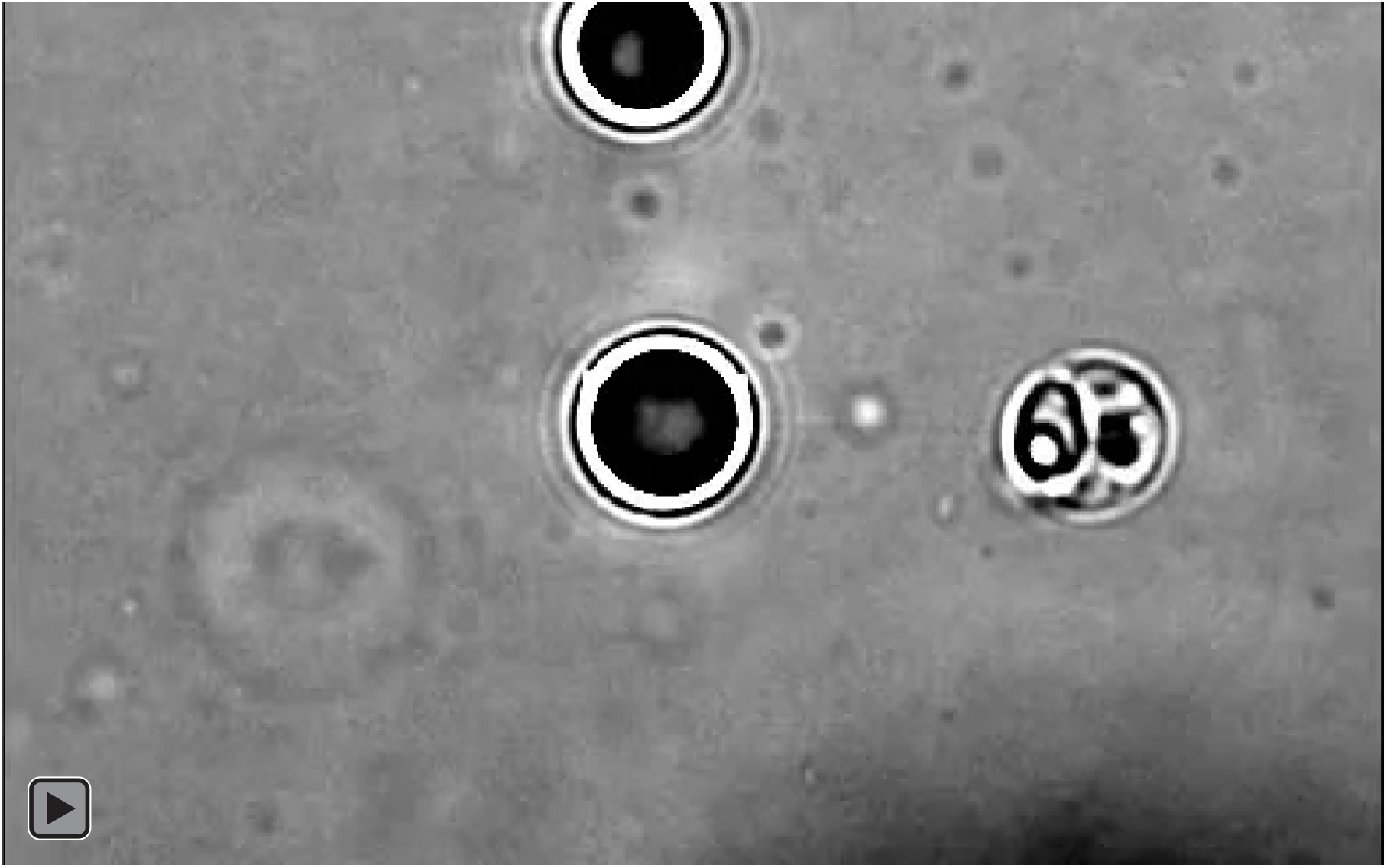
IST1 alone does not promote membrane tube elongation.

**Figure 3.**
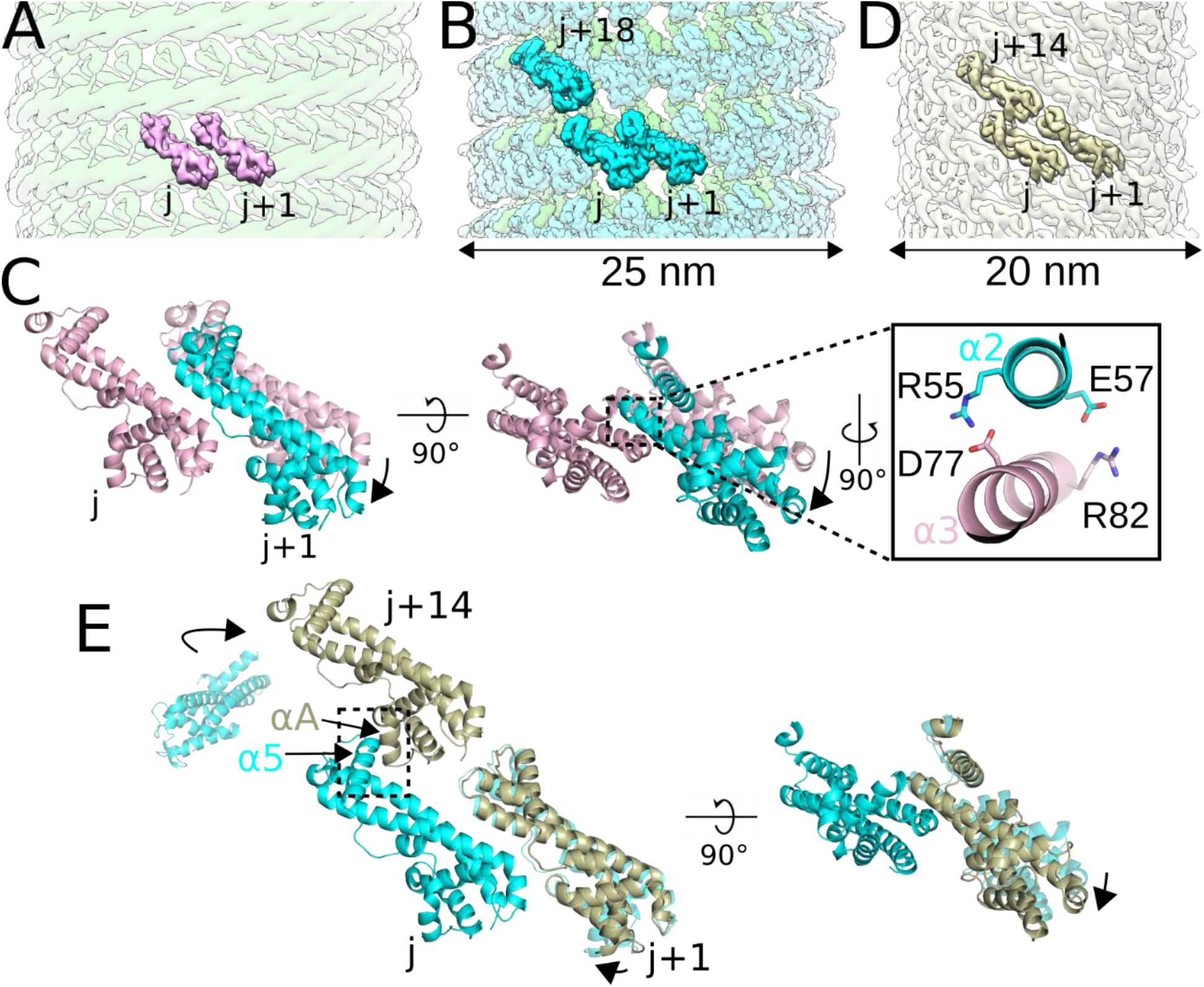
IST1 polymerization drives constriction of the CHMP1B+IST1 filament. (**A**) Model of two IST1 subunits (j, j+1), colored pink, initially binding onto the CHMP1B filament (green). (**B**) CryoEM reconstruction of the right-handed CHMP1B+IST1 filament, with CHMP1B and IST1 in green and cyan, respectively. Three IST1 subunits (j, j+1, j+18) are highlighted.(**C**) Superposition of the j and j+1 IST1 subunits from (A) and (B), with the j subunits used for alignment. (*Inset*), new electrostatic interactions between helix α3 (pink) from the j subunit and helix α2 (cyan) from the j+1 subunit help stabilize intra-IST1 contacts to drive constriction. (**D**) CryoEM 3D reconstruction of the IST1_NTD_^R16E/K27E^ filament (bronze). Three IST1 subunits (j, j+1, j+14) are highlighted. (**E**) Superposition of the j, j+1, and j+14 subunits from the IST1_NTD_^R16E/K27E^ filament in (D) with IST1 subunits from the CHMP1B+IST1 filament in (B). Protomers were aligned by the j subunit. The boxed area highlights helix α5 of the j subunit and helix αA from the j+14 subunit driving inter-turn interactions.

**Figure 3—figure supplement 1.**
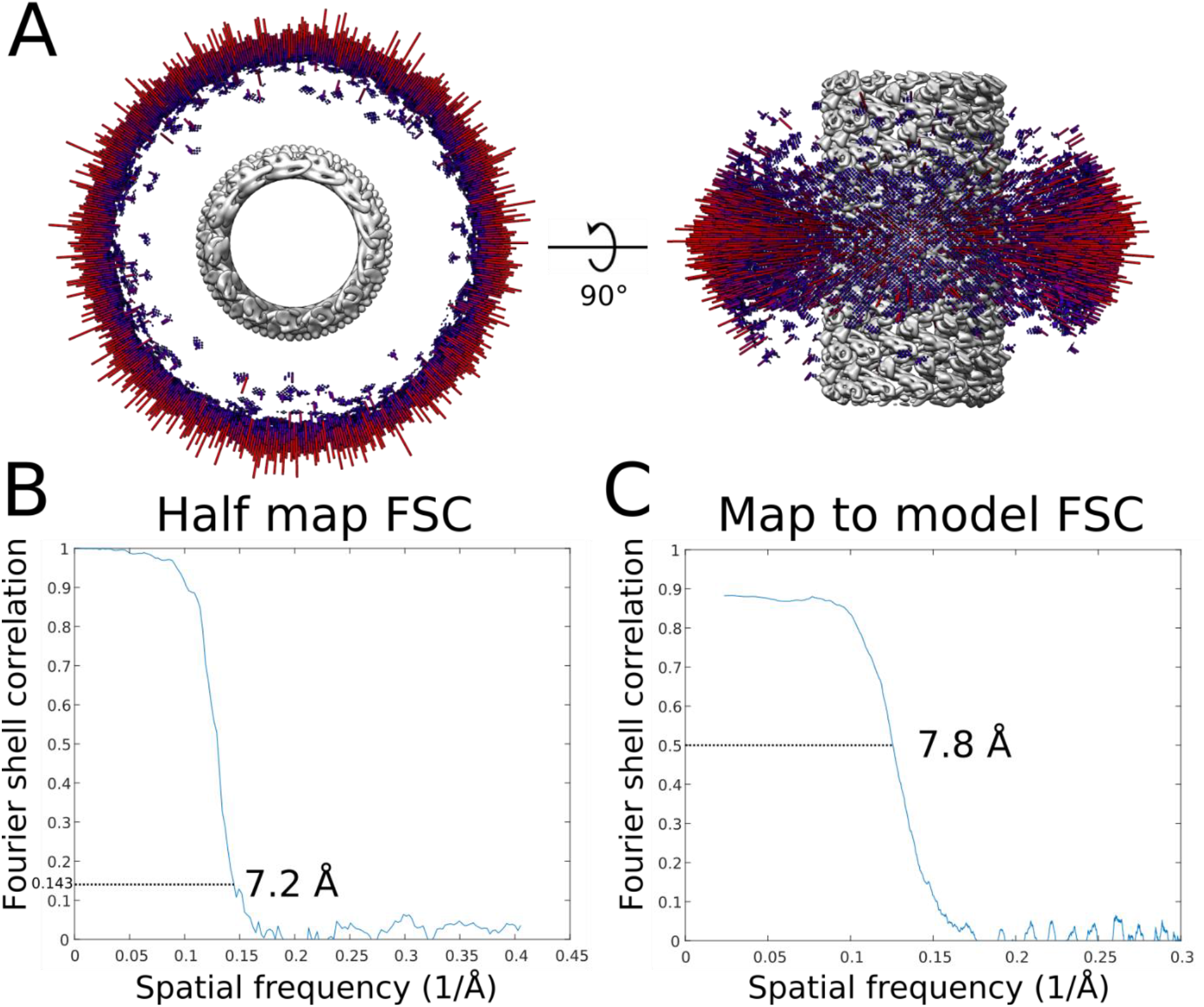
CryoEM validation of the IST1_NTD_^R16E/K27E^ filament. (**A**) Angular distribution of the IST1NTDR16E/K27E filament. (**B**) Half map FSC of the IST1NTDR16E/K27E filament. (**C**) Map to model FSC of the IST1NTDR16E/K27E filament.

**Figure 3—figure supplement 2.**
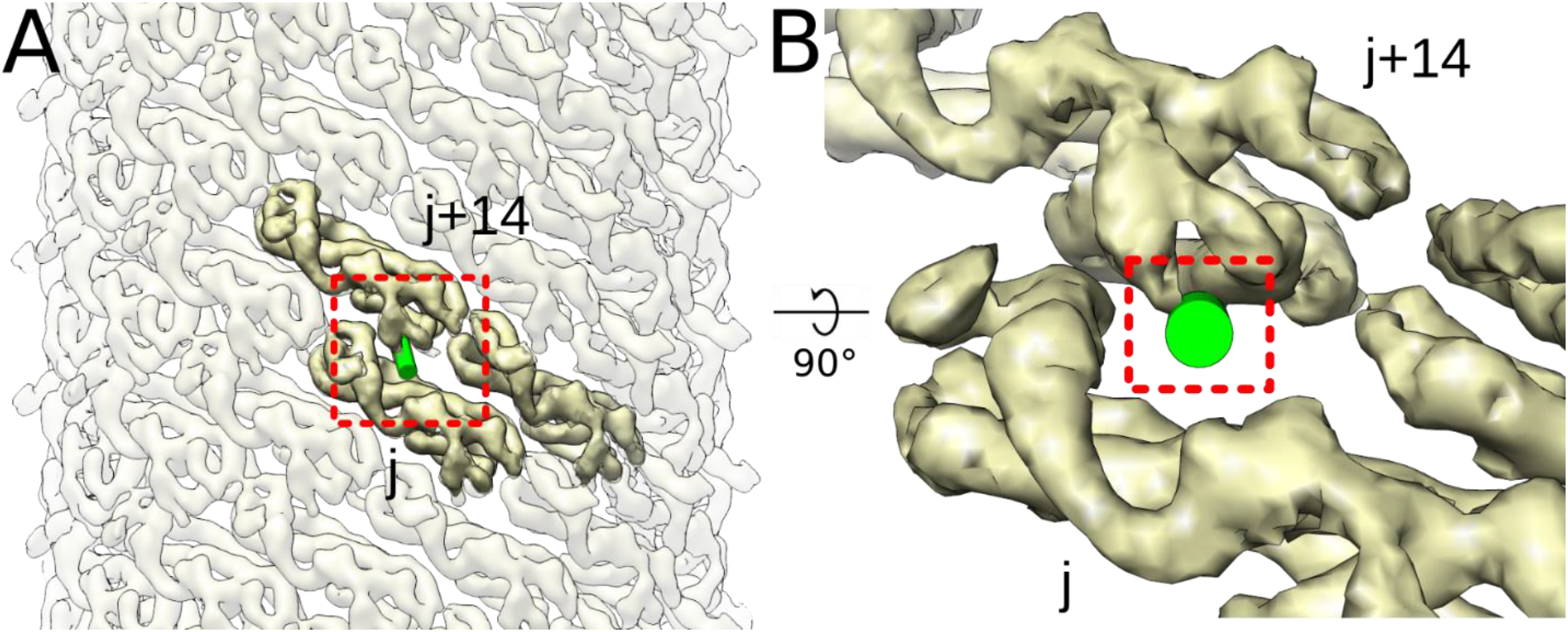
Steric clashing between the CHMP1B MIM and inter-turn IST1 subunits would prevent IST1 from achieving its preferred curvature in the copolymer. (**A**) External view of the IST1_NTD_^R16E/K27E^ filament with one CHMP1B MIM (shown as a green cylinder) docked onto the IST1_NTD_^R16E/K27E^ j subunit. (**B**) Zoomed in view of boxed area in (A) highlighting how the CHMP1B MIM clashes with the IST1 j+14 subunit.

**Figure 3—movie supplement 1.**
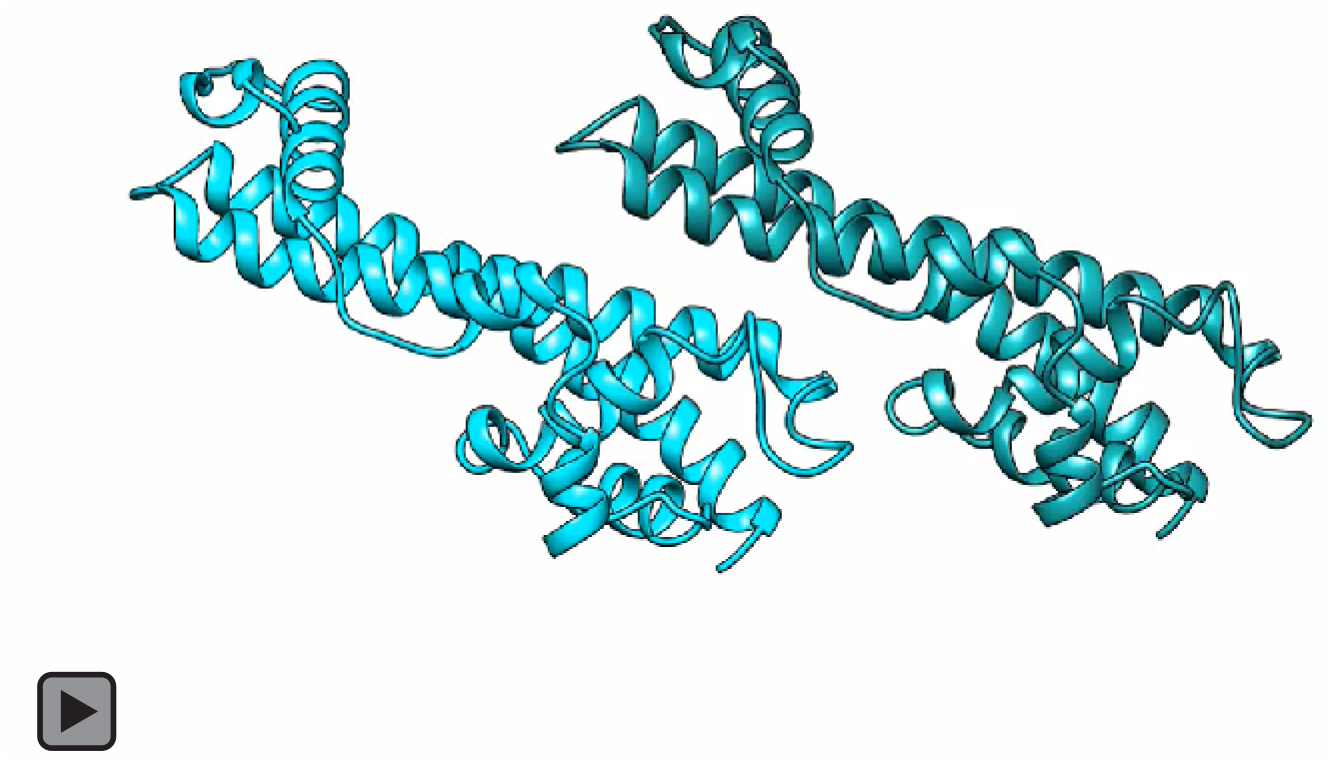
Swinging of two IST1 subunits from initial binding to the CHMP1B filament to the constricted state (side view).

**Figure 3—movie supplement 2.**
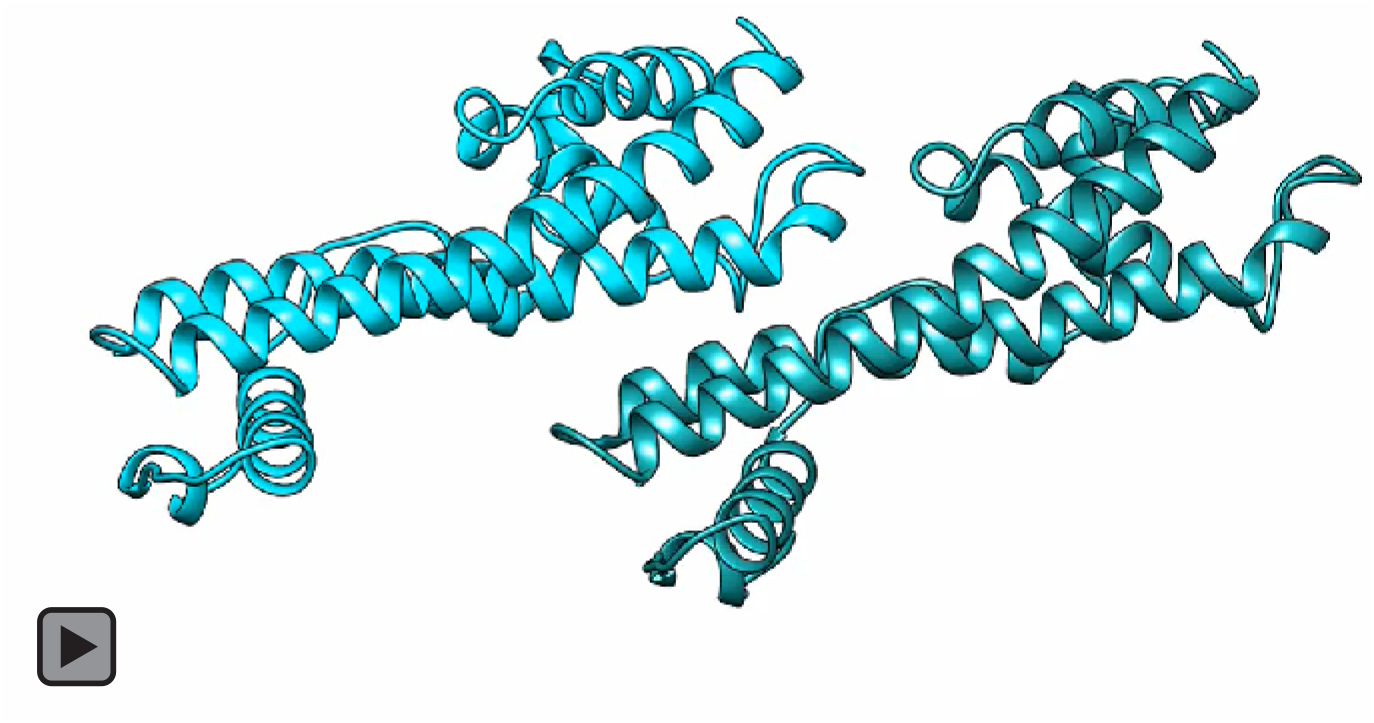
Swinging of two IST1 subunits from initial binding to the CHMP1B filament to the constricted state (top-down view). The membrane would lie at the top of the animation.

The left-handed helical filament of the membrane-bound CHMP1B+IST1 copolymer also a one-start, double-stranded filament. The left-handed copolymer is slightly more constricted, with an outer diameter of 24 nm and an inner leaflet distance of 4.4 nm (Figure 1—figure supplement 2A).

Previous work has documented that ESCRT-III filaments (*30*), like bacterial flagella (*31, 32*), can adopt both left- and right-handed helical structures in vivo. To understand the structural basis of this flexibility, we compared CHMP1B protomer conformations within the left-versus right-handed copolymers (Figure 1—figure supplement 2B–2C). The overall root mean square deviation of the Cɑ backbone (RMSD) between helices ɑ1-ɑ5 of CHMP1B protomers in the two filaments is 1.2 Å. However, the RMSD is smaller when only aligning either the N-terminal helices ɑ1-ɑ2 (∼0.7 Å) or the C-terminal helices ɑ4-ɑ5 (∼0.6 Å) (Figure 1—figure supplement 2B–2C). Therefore, switching between a left-or right-handed filament can be achieved simply by a small change in the joint between helices ɑ2 and ɑ3 (Figure 1—figure supplement 2B–2C).

Sequential assembly of the IST1 strand does not discriminate between the left- and right-handed filaments. The RMSD of single IST1 protomers or between “j” and “j+1” subunits between the two copolymers are only 0.5 Å and 0.6 Å respectively. There are no significant differences in how IST1 assembles around either the left- or right-handed filaments. We envision that as one turn of CHMP1B finishes a revolution, it will either continue polymerizing ‘above’ or ‘below’ the initiating subunit, and this choice will define the helical hand. Thus, it appears that a stochastic flexing between helices ɑ2-ɑ3 will allow either handedness to propagate. We did not observe left-handed helical polymers in our prior work on the lipid-free or the nucleic-acid templated copolymer structure, perhaps due to the different solution and nucleation conditions that promoted lipid-free polymerization (*14, 29*). While we did not reconstruct a left-handed membrane-bound CHMP1B-only filament, this likely simply reflects the limited dataset size and diameter variability.

### The joint between CHMP1B helices ɑ3 and ɑ4 allows for different curvatures

To understand how CHMP1B polymers adopt different curvatures, we examined how the CHMP1B conformation changes between the “moderate” constriction CHMP1B-only and the “high” constriction CHMP1B+IST1 filaments. In all reconstructions, a single CHMP1B protomer (“i”) interacts with eight other protomers in an interconnected network. Among these, helix ɑ5 of the “i” subunit passes behind three neighboring subunits and binds the closed end of the ɑ1-ɑ2 hairpin of the “i+4” subunit (Figure 2A). Thus, CHMP1B subunits always interweave with the same protomers, regardless of the filament’s overall degree of curvature. There is little contact between neighboring turns of the CHMP1B filament in either state, suggesting that turns can slide past one another during constriction (Figure 2D, Figure 2—movie supplement 1, Figure 2—movie supplement 2).

We next compared the conformation of a CHMP1B protomer between the CHMP1B-only and the right-handed CHMP1B+IST1 membrane-bound structures (Figure 2C). The RMSD of helices ɑ1-ɑ5 is 2.4 Å between the two states. However, the RMSD drops to 1.1 Å when comparing only the N-terminal helices ɑ1-ɑ2. Similarly, comparing just the C-terminal helices ɑ4-ɑ5 lowers the RMSD to 1.5 Å. While there is again a small change in the angle between helices ɑ2-ɑ3, the largest conformation difference is at the “elbow” between helices ɑ3-ɑ4 (Figure 2B-C). This flex in the elbow joint, when propagated across an entire turn of the helical assembly, leads to membrane constriction and also tubule elongation (Figure 2D, Figure 2—movie supplement 1, Figure 2—movie supplement 2). To confirm this tube elongation, we used holographic optical tweezers to hold traptavidin-coated beads and pull membrane tubes from giant unilamellar vesicles (GUVs), and then visualized changes in membrane tube lengths as a function of ESCRT-IIIs. CHMP1B addition elongated the tubes slightly, and subsequent incorporation of IST1 produced even longer tubes (Figure 2—figure supplement 1, Figure 2—movie supplement 3). In agreement with our structural studies, tube elongation was dependent on CHMP1B as addition of IST1 alone did not induce membrane elongation (Figure 2—movie supplement 4).

**Figure 4.**
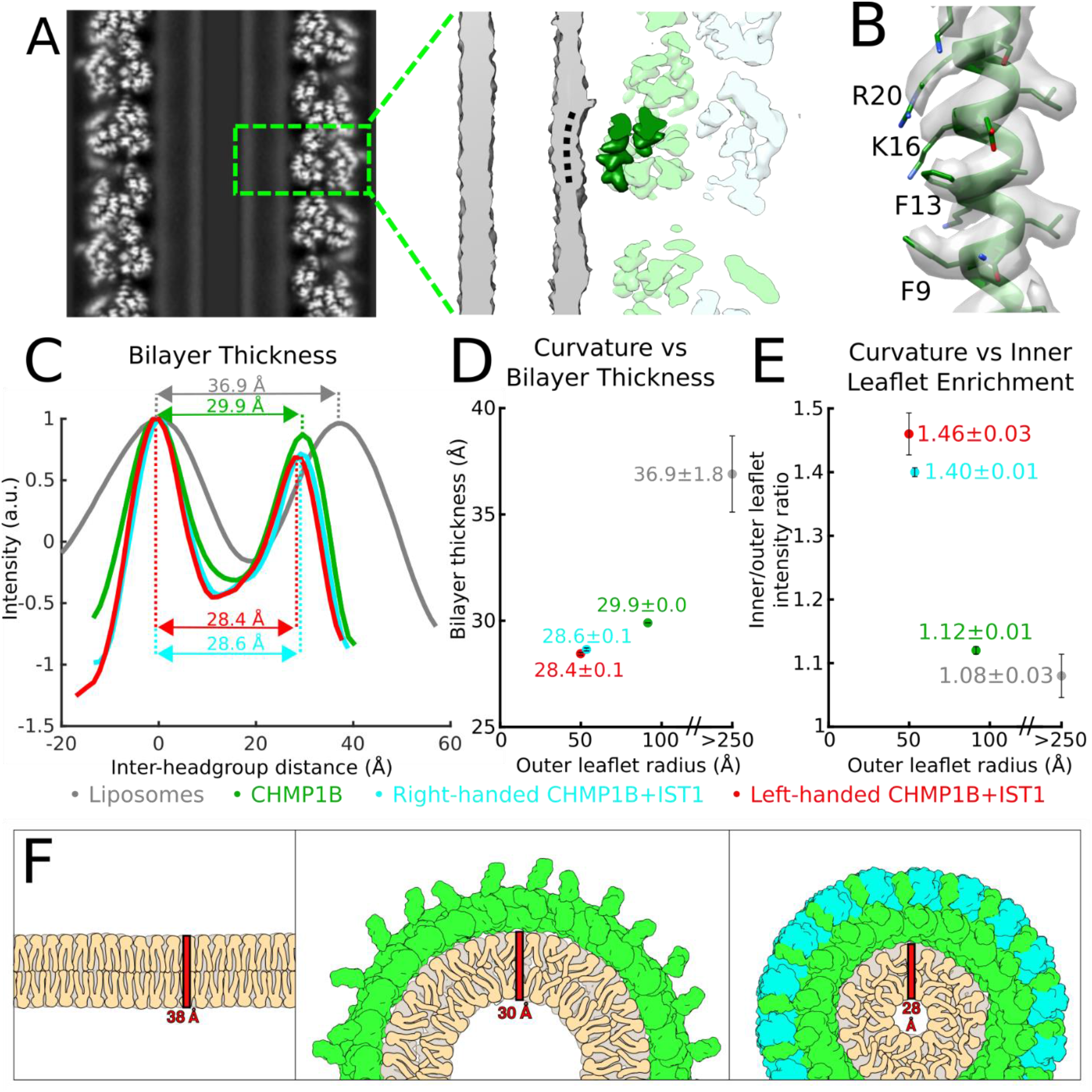
The bilayer thins and the inner leaflet crowds under high curvature. (**A**) The outer leaflet buckles under extreme curvature. *Left*, central slice along the helical axis. *Right*, zoomed view of boxed area in left showing CHMP1B dimpling (black dashed curved line) in the outer leaflet of the bilayer. A CHMP1B helix α1 sitting at the membrane is highlighted in dark green. (**B**) CryoEM density map and model of CHMP1B helix α1 from (A). The residues involved in membrane binding are labeled. (**C**) Intensity plots of membrane bilayer thickness for liposomes only (from 2D class averages, colored in grey), membrane-bound CHMP1B (green), and right-or left-handed membrane-bound CHMP1B+IST1 filaments (cyan and red respectively) as determined by cryoEM. The bilayer thickness is labeled for each. Intensities were normalized to the peak intensity of the inner leaflet. (**D**) Plot of membrane thickness as a function of radius of the outer leaflet. The dots are colored as in (left). For (C) and (D), the liposome values were determined from 2D averages (n=6) while the others were determined from half maps from each 3D reconstruction (n=2). (**E**) The inner membrane density increases as a function of curvature. Plot of ratio of inner leaflet to outer leaflet peak intensity as a function of radius of the outer leaflet. Dots are labeled as in (C). (**F**) Schematic illustration of lipid behavior as the bilayer is remodeled from planar (*left*), to moderate curvature by CHMP1B (*middle*), to high curvature by CHMP1B+IST1 (*right*). The outer leaflet headgroups spread out, while the inner leaflet headgroups crowd and the aliphatic tails become more disordered and therefore less extended.

**Figure 4—figure supplement 1.**
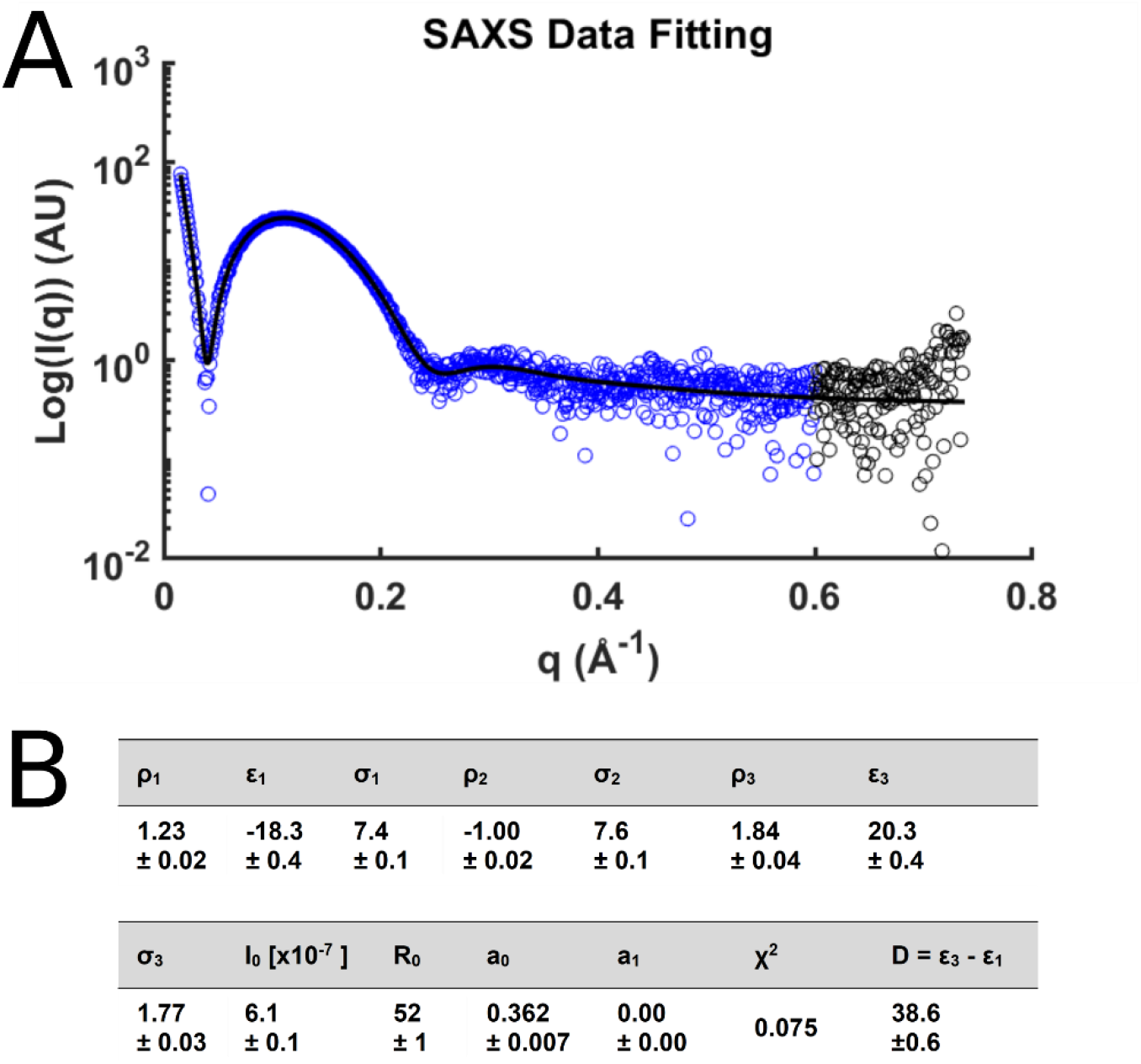
SAXS analysis of liposomes and calculation of bilayer thickness. (**A**) The small angle scattering intensities for protein-free unilamellar vesicles used in this study. The black line represents the fit to the model. The blue data points were used for fitting. (**B**) Fit results for the liposomes and the resulting thickness, D (Å). The bilayer center, ε_2_, was fixed at 0, and the magnitude of the central peak, ρ_2_, was fixed at −1.

**Video 1.**
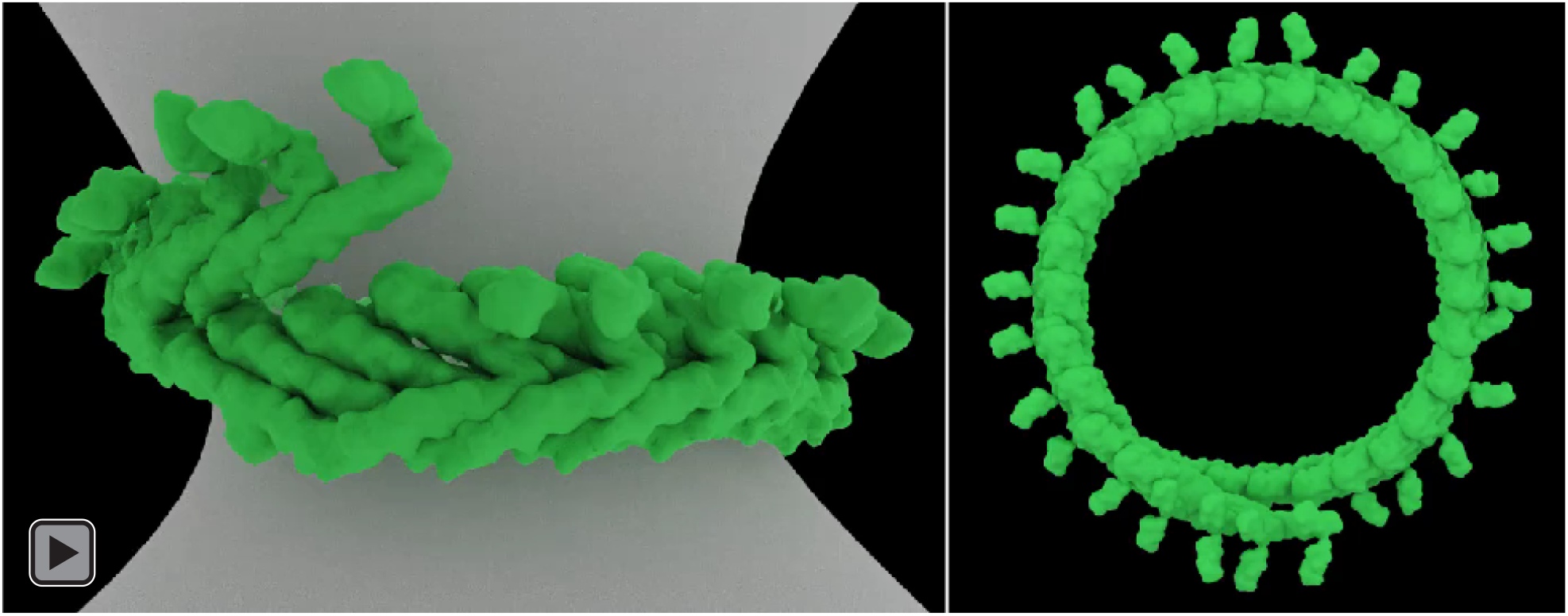
Constriction of a membrane tubule by CHMP1B and IST1.

Previous work has shown that other ESCRT-IIIs like CHMP2, CHMP3, and CHMP4 also form filaments with a wide range of curvatures (*33–35*). Owing to the high homology of the ESCRT-III core, we suggest that these proteins also have dynamic elbow joints that will accommodate changes in filament curvature. As noted above, there are minimal contacts between turns of CHMP1B to stabilize inter-turn interactions (Figure 2—movie supplement 2). Indeed, in vivo images of ESCRT-IIIs at different sites of action reveal conical spirals with significant gaps between turns (*35–37*). Thus, the inherent flexibility of ESCRT-III subunits could allow such loosely packed filaments to form with a range of diameters and helical pitches, and to slide past one another upon constriction.

### IST1 polymerization drives constriction

To understand how IST1 induces constriction of CHMP1B-membrane filaments, we first docked two IST1 subunits (j and j+1) onto sequential CHMP1B protomers (i and i+1) of the moderate-curvature CHMP1B-only membrane, based on the CHMP1B-IST1 intersubunit interactions observed in the copolymer filaments (Figure 3A). We then compared how the interactions between IST1 subunits would change between this ‘initial IST1 binding’ state and those observed in the CHMP1B+IST1 filaments (Figure 3B–3C). The IST1 j+1 subunit from the copolymer subunit swings closer (∼8 Å) to the j subunit when compared to the j+1 subunit from the initial IST1 binding state, adding 480 Å^2^ of buried surface area (BSA) (Figure 3—movie supplement 1, Figure 3—movie supplement 2). This swing enables the formation of hydrogen bonds between IST1 residues D77 and R82 on helix ɑ3 of the j subunit with R55 and E57 on helix ɑ2 of the j+1 subunit (Figure 3C). In contrast, IST1 subunits make only minimal contacts between adjacent turns (15 Å^2^ of BSA between the j and j+18 subunits and no contacts between the j+1 and j+18 subunits). Thus, interactions between the j and j+1 subunits along the IST1 strand provide the force that flexes the CHMP1B elbow and consequently constricts and elongates the filament. These sliding, lateral interactions to promote changes in filament architecture have also been observed for the yeast ESCRT-III proteins Snf7 and Vps24 (*38*).

IST1 has recently been reported to induce constriction of ESCRT-III assemblies in vitro (*39*). To understand how IST1 polymerization could drive constriction, we determined a 3D reconstruction of a protein-only IST1 filament (Figure 3D). We previously showed that an N-terminal construct of IST1 (residues 1-189) harboring R16E and K27E mutations (IST1_NTD_^R16E/K27E^) cannot co-assemble with CHMP1B as the mutations destabilize the CHMP1B-IST1 interface, but can still form tightly-packed homopolymeric helical tubes (*14*). The IST1-only filaments were heterogeneous in diameter (∼18-28 nm), with the majority of filaments narrower than the CHMP1B+IST1 copolymers (24-25 nm). We identified a major subset of IST1_NTD_^R16E/K27E^ tubes that were ∼20 nm wide and reconstructed this class to moderate resolution, revealing the secondary structure elements of closed-conformation IST1_NTD_ subunits (Figure 3—figure supplement 1). IST1_NTD_^R16E/K27E^ forms a single-stranded, right-handed filament (Figure 3D). In the absence of a membrane or CHMP1B, therefore, IST1 alone polymerizes into even narrower helical assembly than either of the copolymers. Interestingly, the reconstruction of these IST1-only filaments revealed that IST1 always adopted the closed conformation, suggesting that IST1 may exclusively function in the closed state.

To understand how the arrangement of IST1 subunits between IST1-only and CHMP1B+IST1 filaments differ, we compared the interactions of IST1 subunits at the inter-turn interface between the copolymer (j, j+1, and j+18 subunits) and the IST1_NTD_^R16E/K27E^ (j, j+1, and j+14 subunits) filaments. The RMSD between a protomer from the IST1_NTD_^R16E/K27E^ filament and an IST1 protomer from the copolymer filament is ∼1 Å, with minimal changes in the Cα backbone. The j and j+1 contact that defines nearest-neighbor IST1-IST1 interactions is conserved in both structures, but the IST1 j and j+1 subunits swing slightly closer together (∼2 Å) in the IST1_NTD_^R16E/K27E^ filaments (Figure 3E). Propagation of this subtle change actually decreases the BSA between the j and j+1 subunits (from 480 Å^2^ to 340 Å^2^), but increases inter-turn contacts, which are predominantly made by helix α5 of the j+1 subunit contacting helix αA of the j+14 subunit (220 Å^2^) and additional packing between the j+1 and j+14 subunits (100 Å^2^). This results in an overall increase of 165 Å^2^ of BSA at the inter-turn interface for the IST1_NTD_^R16E/K27E^ filament (Figure 3E). Thus, unlike the constriction seen upon addition of IST1 to the CHMP1B filament, this second constriction step appears to be driven by inter-turn contacts.

We note that the inter-turn interactions involving the j and j+14 subunit in the IST1-only filament are unattainable in the CHMP1B+IST1 filament, as the presence of the CHMP1B helix α6, the MIT interacting motif (MIM), sterically blocks the IST1 j and j+14 packing (Figure 3—figure supplement 2). Intriguingly, the interaction between IST1 helix α5 and the CHMP1B MIM supports efficient assembly of the copolymer (*29*). We speculate that modulation of this interface by the VPS4 family of ATPases could, in principle, regulate the degree of constriction achieved by the CHMP1B+IST1 filament.

### The membrane bilayer thins and the inner leaflet compresses at high curvature

We also examined the consequences to the membrane as a function of increasing curvature stress (Figure 4). In both copolymer reconstructions, helix ɑ1 of CHMP1B faced the membrane, and the surface of the bilayer appeared to be “dimpled” by certain amino acids (Figure 4A). Specifically, the conserved CHMP1B residues F9, F13, K16, and R20, which lie on the same face of helix ɑ1, comprise the protein-membrane interface (Figure 4B). No residues appeared to insert deeply into the membrane. Rather, they appear to sit at the hydrated surface of the bilayer. The mixed aromatic and cationic character of the CHMP1B region that most closely approaches the membrane does not suggest any lipid recognition specificity beyond net anionic charge, with the basic residues complementing the negatively charged membrane. It is somewhat surprising that the hydrophobic residues remain fully hydrated at this degree of constriction, but we speculate that they may be poised to insert into the membrane (see below).

By measuring the peak-to-peak distances between the outer leaflet and inner leaflet headgroups, we observed a correlation between bilayer thinning and the degree of membrane constriction by CHMP1B and by CHMP1B+IST1. To determine the bilayer thickness of our initial, unconstricted bilayers, we performed small-angle X-ray scattering (SAXS) of our relaxed liposomes. This experiment yielded a thickness estimate of 38.6 ± 0.6 Å (Figure 4—figure supplement 1), consistent with previously published SAXS measurements of membranes composed primarily of SDPC (*40*). We compared this measurement with bilayer profiles from cryoEM 2D averages of segments of our liposomes, which yielded a thickness of 36.9 ± 1.8 Å, which is within the experimental uncertainty of the SAXS measurement (Figure 4C–4D). Upon constriction by CHMP1B alone, the bilayer compressed to 29.9 ± 0.0 Å. The sequential addition of IST1 led to further compression of the membrane to 28.6 ± 0.1 Å and 28.4 ± 0.1 Å for the right-handed and left-handed copolymers, respectively (Figure 4C–4D). In addition, the intensity of the inner leaflet increased as a function of constriction, with inner/outer leaflet peak intensities of 1.08 ± 0.03, 1.12 ± 0.01, 1.40 ± 0.01, and 1.46 ± 0.03 for the liposomes, moderate-constriction CHMP1B filaments, and the high-constriction right-handed and left-handed CHMP1B+IST1 filaments, respectively (Figure 4E). Thus, the lipid density in the inner leaflet increases significantly as the membrane tubule constricts towards the fission point.

In agreement with physical models of membrane behavior (*41, 42*), our reconstructions indicate that the bilayer thins as the membrane is constricted (Figure 4C–4D) and that the outer leaflet headgroups separate while the inner leaflet headgroups become more crowded (Figure 4E). To accommodate this thinning, the acyl chains from both leaflets likely become more disordered and less extended (Figure 4F). It has also been suggested that curvature stress causes local lipid composition changes that increases the membrane line tension and promote fission (*43, 44*). We speculate that these changes may lower the activation barrier for fission. With this specific lipid composition, however, the tubes remain stable with only a 4.2 nm gap between the inner leaflet headgroups across the lumen.

## Conclusion and Perspective

A remaining central question is how ESCRT-III proteins work with their associated AAA ATPases to catalyze membrane constriction and, ultimately, membrane fission (*12, 33, 34, 45-52*). Recent in vivo and in vitro studies have explored the roles of staged ESCRT-III assembly (*39*) and ATP-dependent forces (*39, 53*). Here, we show how CHMP1B and IST1 function sequentially to squeeze and thin the membrane, bringing it nearly to the fission point (Video 1). Specifically, CHMP1B first binds and assembles into a flexible filament that wraps the target into a moderate curvature tubule. CHMP1B flexibility allows for many degrees of filament curvature and handedness, which may explain how it and other ESCRT-III proteins can adopt a wide range of architectures. IST1 binding then drives CHMP1B to constrict the membrane even further, exploiting IST1-intersubunit interactions to form tighter and tighter turns. Lipid composition is also expected to play a role in this process, although our work does not address that issue directly.

Finally, our reconstructions suggest that IST1 may drive CHMP1B into an even narrower constriction state during the fission step, and that this process could be regulated by a VPS4 family member. Comparison of the CHMP1B+IST1 and IST1-only filaments suggests that CHMP1B helix α6 (MIM) may sterically limit the full potential of IST1 constriction. We therefore speculate that unfolding or displacement of the CHMP1B MIM could trigger further tightening of the double-stranded filament. Importantly, the CHMP1B MIM forms the binding site for the MIT domains of VPS4 ATPase family members (*54–56*), and MIT domain binding could therefore provide a mechanism for displacing this helix (*34, 35, 39, 48*). Further constriction of the CHMP1B+IST1 filament might also push the two aromatic residues of CHMP1B helix α1 (F9 and F13) from the hydrated surface layer, as seen in our structure, into the hydrophobic interior of the outer membrane leaflet, thereby destabilizing the membrane, helping to drive lipid mixing, and promoting fission (*57, 58*).

## Materials and Methods

### Protein Expression and Purification

The purification of the N-terminal domain of IST1 containing residues 1-189 (IST1_NTD_) and the N-terminal IST1 R16E/K27E mutant (IST1_NTD_^R16E/K27E^) have been described previously (*14*). CHMP1B residues 1-199 were cloned into an N-terminal 6xHis-SUMO fusion to yield a native N-terminus after removal of the purification tag. Two alleles of CHMP1B have been reported, 37K or 37E, and we saw that both alleles remodeled membranes and copolymerized with IST1 with indistinguishable activity. The CHMP1B 37E allele was used for subsequent studies and was expressed in LOBSTR-BL21 (DE3) cells (*59*) in ZYP-5052 auto-induction media (*60*). Cells were harvested and frozen at −80 °C. All subsequent steps were performed at 4 °C unless otherwise noted. Thawed cells were suspended in lysis buffer (50 mM Tris, pH 8, 500 mM NaCl, 10 mM Imidazole, 1 mM DTT, 5% (v/v) glycerol) and supplemented with lysozyme. Cells were lysed by sonication. Lysate was centrifuged at 30,000 x g for 1 h, and the supernatant was filtered using a 0.45 µm membrane. Clarified lysate was loaded onto a gravity flow column with Ni-NTA resin (Qiagen), incubated for 1 hour, and washed extensively with lysis buffer. The fusion protein was eluted in lysis buffer supplemented with 400 mM imidazole. His-tagged ULP1 protease was added and then dialyzed into cleavage buffer (20 mM Tris pH 8.0, 150 mM NaCl, 1 mM DTT, 10 mM imidazole) at room temperature overnight. The cleaved product was then applied again to Ni-NTA resin to remove the purification tag, uncleaved fusion protein, and the protease. CHMP1B was further purified by Superdex-75 16/60 size exclusion chromatography (GE Healthcare Life Sciences, USA) in size exclusion buffer (20 mM Tris, pH 7.4, 150 mM NaCl, 1 mM DTT).

### Liposome Preparation

Stock lipid solutions (Avanti Polar Lipids) were resuspended in chloroform. To produce the liposomes, (18 mole% 16:0-18:1 phosphatidylserine (POPS), 58% 16:0-22:6 phosphatidylcholine (SDPC), 18% cholesterol, 6% PI(3, 5)P_2_ or equivalent phosphoinositide), 2 mg total lipid were dried in a glass vial at room temperature under streaming nitrogen with vortexing. The lipids were again re-dissolved in chloroform, dried under streaming nitrogen, and desiccated under house vacuum (at least 4 hours in darkness). The lipid films were dispersed in 1 ml buffer (25 mM Tris, pH 7.4, 150 mM NaCl, 2 mg/ml final concentration, 4 °C overnight with gentle rocking). Liposomes were freeze-thawed 10 times and then stored at −80 °C.

### Membrane Remodeling Reactions

Membrane remodeling reactions were performed at room temperature with protein concentrations ranging from 5 – 15 µM and liposome concentrations ranging from 0.5 – 1 mg/ml in reaction buffer (20 mM Tris, pH 7.4, 150 mM NaCl). CHMP1B was incubated with liposomes overnight at room temperature. For membrane-bound CHMP1B reactions, these were then directly used for EM sample preparation. For samples including IST1, the sample was pelleted (13,000 x g, 5 mins), the supernatant was decanted to remove unbound CHMP1B, and the pellet was resuspended in reaction buffer with equimolar IST1_NTD_. This was incubated for 10 mins and then subjected to EM sample preparation.

### CryoEM Sample Preparation and Data Collection

3.5 µL of the membrane remodeling reactions were applied to glow-discharged R1.2/1.3 Quantifoil 200 Cu mesh grids (Quantifoil) in a Mark III Vitrobot (FEI). Grids were blotted with Whatman #1 filter paper (Whatman) for 4-8 seconds with a 0 mm offset at 19 °C and 100 % humidity before plunging into liquid ethane. Grids were stored under liquid nitrogen until samples were imaged for structural determination. Datasets were collected either on a 300 kV Technai Polara, a 300 kV Titan Krios, or a 200 kV Technai F20, all using a K2 Summit detector operated in super-resolution mode and binned by a factor of 2 for subsequent processing. Data collection parameters are summarized in Table 1.

### EM Image Analysis and 3D Reconstructions

All dose-fractionated image stacks were corrected for motion artifacts, 2x binned in the Fourier domain, and dose-weighted using MotionCor2 (*61*). GCTF-v1.06 (*62*) was used for contrast transfer function (CTF) estimation. Particles were selected manually using RELION3 with helical processing (*63*), and subsequent steps were performed in RELION3 unless otherwise stated. Segments were extracted with ∼90% overlap between boxes. Multiple rounds of 2D classification were performed to remove poor particles and to yield particles with a largely uniform diameter. For determination of the helical parameters for the CHMP1B-only or IST1_NTD_^R16E/K27E^ filaments, the iterative helical real space refinement (IHRSR) algorithm (*64*) as implemented in SPIDER (*65*) was used. For the CHMP1B+IST1 filaments, the previously determined helical parameters were used as the initial values (*14*). For all reconstructions, hollow, smooth cylinders were used as initial models for 3D auto-refine reconstructions with refinement of helical parameters and a central Z length of 40% of the particle box and the ‘ignore CTFs until first peak’ flag for CTF estimation was used. These particles then went through multiple rounds of 3D classification without alignment with 3-4 classes and T values varying from 2-10 and a protein-membrane mask. Selected particles then were subject to another 3D auto-refine reconstruction with per-particle CTF estimation correction within RELION3, followed by a final 3D auto-refine reconstruction. For the membrane-bound left-handed CHMP1B+IST1 reconstruction, initial 3D classification without alignment yielded a class with no discernable features. 2D classification of these particles still yielded class averages with secondary structure features. Refinement of the helical parameters by switching the sign of the twist was then able to generate a good initial 3D reconstruction that was further refined as above. Post-processing was performed using the masks from refinement with ad hoc B-factors applied. Refinement parameters are listed in Table 2.

### Atomic Modeling and Validation

For the high-resolution CHMP1B+IST1 filaments, a single protomer of CHMP1B and IST1 from the previously determined structure were initially docked into the density with UCSF Chimera (*66*). The protomers were manually adjusted and rebuilt in *Coot* (*67*) and then refined in phenix.real_space_refine (*68*) using global minimization, morphing, secondary structure restraints, and local grid search. The refined protomers were then used to generate roughly two full turns manually in real space using UCSF Chimera. Noncrystallographic symmetry (NCS) constraints were then used thorough refinement in phenix.real_space_refine. Iterative cycles of manually rebuilding in *Coot* and phenix.real_space_refine, with previous strategies and additionally B-factor refinement, were performed. For the low resolution CHMP1B-only or IST1_NTD_ filaments, a similar procedure was performed but only roughly one turn was built and no B-factor refinement was performed in phenix.real_space_refine.

All final model statistics were tabulated using Molprobity (*69*) (Table 3). Map versus atomic model FSC plots were computed in PHENIX (*70*). All structural figures were generated with UCSF Chimera and PyMOL (*71*).

### Real-time Imaging of Membrane Constriction

Lipid tube pulling and imaging were performed using a previously described holographic optical trapping setup possessing an independent fluorescent imaging capability (*72*). Briefly, the setup included a custom-modified Eclipse Ti microscope (Nikon Instruments; Melville, NY), with a Nikon 100X, 1.49NA oil immersion objective, Sapphire 488 excitation laser (Coherent, Santa Clara, CA), and a DU897 camera (Andor Technology, Oxford Instruments, USA). All videos were recorded at 20 fps. Biotinylated giant unilamellar vesicles (GUVs) in Flow Buffer (50 mM Tris pH 8.0, 150 mM NaCl, 1 mM DTT,) were synthesized as previously described (*73*). Biotinylated silica beads (5 μm diameter, Si5u-BN-1, Nanocs, USA) were incubated with 10 μM traptavidin (Kerafast, USA) in PMEE buffer (35 mM PIPES, 5 mM MgSO4, 1 mM EGTA, 0.5 mM EDTA, pH 7.2) for 30 min followed by two rounds of washing (centrifugation at 12,000 × g for 2 minutes followed by supernatant removal and re-suspension of the pellet in 50 mM Tris buffer) to remove excess traptavidin. The beads were then mixed with GUVs and diluted as necessary to limit bead density to 1-3 beads in each 50 μm x 50 μm field of view. The mixture was then applied to a flow cell and immediately subjected to experiments.

Membrane tubules were formed by capturing a bead in an optical trap, moving the trapped bead into contact with a GUV, and finally, upon attachment, moving the bead away from the GUV. The presence of a tubule was then assessed visually in real time. The GUVs were stabilized in place via other beads attached to their surface and held in independent holographically defined traps or via non-specific surface attachment. Proteins were introduced into the flow cell in a sequential manner: first 0.5 μM CHMP1B and then 0.5 μM IST1 (both in Flow Buffer). Lipids tubules were held perpendicular to flow direction during these buffer exchanges and the resulting tubule constrictions were recorded. Experiments with CHMP1B and IST1 in slightly different buffers (0.5 μM CHMP1B in 10 mM Tris pH 8.0, 100 mM NaCl 1 mM DTT or 0.5 μM IST1 in 50 mM Tris pH7.0, 350 mM NaCl, 5% (v/v) glycerol, 5 mM 2-mercaptoethanol) yielded similar results.

### Small Angle X-ray Scattering

For small-angle X-ray scattering (SAXS) analysis, liposomes were formed by vortexing a dry lipid film (similar to liposome preparation as stated above) in water to yield a final lipid concentration of 15 mg/ml. Liposomes were extruded through a 100 nm-pore polycarbonate membrane, followed by extrusion through a 50 nm-pore polycarbonate membrane. Synchrotron SAXS data were collected at beamline 4-2 of the Stanford Synchrotron Radiation Lightsource (SSRL), Menlo Park, CA (*74*). The sample to detector distance was set to 1.1 m, and the X-ray wavelength used was λ = 1.127 Å (11 keV). Using a Pilatus 3 × 1M detector (Dectris Ltd, Switzerland) the setup covered a range of momentum transfer of q ≈ 0.017 – 1.17 Å^-1^ where q is the magnitude of the scattering vector, defined as q = 4π sinθ /λ, where θ is the scattering angle, and λ is the wavelength of the X-rays. Aliquots of 32 μl of freshly extruded vesicles were loaded onto the automated sample loader at the beamline (*75*). Consecutive series of thirty 2 s exposures were collected first from the buffer blank followed by the vesicle samples. Solutions were oscillated in a stationary quartz capillary cell during data collection to reduce the radiation dose per exposed sample volume. The collected data were radially integrated, analyzed for radiation damage and buffer subtracted using the automated data reduction pipeline at the beam line. To improve statistics and check for reproducibility, the measurements were repeated with different aliquots four times. As no significant differences were found between the repeat measurements, the different data sets were averaged together.

The buffer-subtracted and averaged data were fit using a model for unilamellar vesicles (*76*) as previously described (*77*). The electron density profile of the bilayer is approximated by three Gaussian peaks corresponding to the inner and outer phosphate peaks and a negative peak at the center for the hydrocarbon region. The following Equation 1 was used to fit the data:

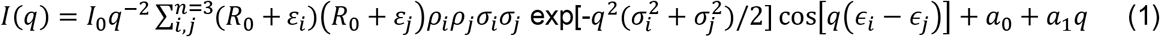

where R_0_ is the mean radius of the vesicle measured from the center of the bilayer, ε is the peak displacement from the bilayer center, σ the Gaussian width of the peak and ρ is its amplitude. I_0_ is the overall intensity of the measured profile. A background term was added, consisting of a constant a_0_ and a linear term a_1_, to take into account the contribution from possible lateral correlations between the lipids. The measured data were fit in the q-region between q = 0.02 –0.6 Å^-1^ (depicted as blue colored points in Figure 4—figure supplement 1A). First, the data were fit using a simulated annealing routine, and the results were then further optimized using a non-linear least square algorithm, both by using code from the open source GNU scientific library project (https://www.gnu.org/software/gsl/). The final fit parameters including the final χ^2^ value of the fits and the resulting bilayer thickness (measured as distance between the inner and outer leaflet peak positions) are summarized in Figure 4—figure supplement 1B.

### Image Analysis of Membrane Bilayer

To determine the bilayer thickness and relative intensity of inner and outer leaflets, Fiji (*78, 79*) was used. The cryoEM half maps from the C1 reconstructions were each low-pass filtered to 8 Å, summed along the central 40% along the Z-axis, and the maps were radially averaged using the Radial Profile Extended plugin (*80*) to determine the intensities as a function of radius, resulting in two measurements per reconstruction. 2D averages of liposomes (n=6) were low pass filtered to 20 Å, parameters for a circle that defined the 2D average were determined, and a wedge that best covered the image was then used to calculate the intensities. For all resulting plots, the radial intensities were normalized to the peak intensity of the inner leaflet peak and then fit to a three Gaussian model (*81*) similar to the SAXS measurements. The model was minimized by a non-linear least square algorithm and the fit errors between the data and models ranged from 1.45%-3.30% with R^2^ values from 0.996-0.999. The local maxima for the headgroups were used to determine the bilayer thickness and inner/outer leaflet intensity ratios.

## Accession Numbers

CryoEM maps and models were deposited to the PDB and EMDB with the following codes: membrane-bound CHMP1B-only filament (PDB ID: 6TZ9, EMD-20590), membrane-bound right handed CHMP1B+IST1 filament (PDB ID: 6TZ4, EMD-20588), membrane-bound left handed CHMP1B+IST1 filament (PDB ID: 6TZ5, EMD-20589), IST1_NTD_^R16E/K27E^ filament (PDB ID: 6TZA, EMD-20591).

## Acknowledgements

We thank members of the Frost lab for helpful discussion, and especially Paul Thomas with computational assistance. We thank Michael Grabe, Maxwell Tucker, David Argudo, Ed Lyman, Alex Sodt, Greg Huber, and Aurelien Roux for discussions on membrane biophysical behavior. We thank Thomas Weiss for assistance with SAXS data collection. We thank Michael Braunfeld, David Bulkley, Matt Harrington, Alexander Myasnikov, and Zanlin Yu of the UCSF Center for Advanced CryoEM for microscopy support. We thank Joshua Baker-LePain and the QB3 shared cluster (NIH grant 1S10OD021596-01) for computational support. Structural biology applications used in this project were compiled and configured by SBGrid (*82*). The Titan X Pascal used for this research was donated by the NVIDIA Corporation. This work was supported by NIH grants P50 082545 and 1DP2GM110772-01 (to A.F.), R01 GM112080 and R37 AI51174 (to W.I.S.), A.F. is further supported by a Faculty Scholar grant from the HHMI and is a Chan Zuckerberg Biohub investigator.

